# Virus-Targeted Transcriptomic Analyses Implicate Ranaviral Interaction with Host Interferon Response in Frog Virus 3-infected Frog Tissues

**DOI:** 10.1101/2021.04.28.441901

**Authors:** Yun Tian, Francisco De Jesús Andino, Jacques Robert, Yongming Sang

## Abstract

Frog Virus 3 (FV3) is a large dsDNA virus that cause global infections in amphibians, fish and reptiles, and contribute to amphibian declines. FV3’s genome contains near 100 putative open reading frames (ORFs). Previous studies have classified these coding genes into temporal classes as immediate early, delayed early and late viral transcripts based on their sequential expression during FV3 infection. To genome-wide characterize ranaviral gene expression, we performed a whole transcriptomic analysis (RNA-Seq) using total RNA samples containing both viral and cellular transcripts from FV3-infected *Xenopus laevis* adult tissues using two FV3 strains, a wild type (FV3-WT) and an ORF64R-deleted recombinant (FV3-Δ64R). In samples from the infected intestine, liver, spleen, lung and especially kidney, a FV3-targeted transcriptomic analysis mapped reads spanning the full-genome coverage at ∼10× depth on both positive and negative strands. By contrast, reads were only mapped to partial genomic regions in samples from the infected thymus, skin and muscle. Extensive analyses validated the expression of almost all annotated 98 ORFs and profiled their differential expression in a tissue-, virus-, and temporal class-dependent manners. Further studies identified several putative ORFs that encode hypothetical proteins containing viral mimicking conserved domains found in host interferon (IFN) regulatory factors (IRFs) and IFN receptors. This study provides the first comprehensive genome-wide viral transcriptome profiling during infection and across multiple amphibian host tissues that will serve as instrumental reference. It also presents evidence implying that ranaviruses like FV3 have acquired previously unknown molecular mimics interfering with host IFN signaling during evolution.

**Importance:** Frog Virus 3 (FV3), are large dsDNA viruses that cause devastating infections globally in amphibians, fish and reptiles, and contribute to catastrophic amphibian declines. FV3’s large genome encodes near 100 coding genes, of which most have been functionally uncharacterized in the viral pathogenesis. Using a whole transcriptomic analysis (RNA-Seq) in FV3-infected amphibian samples, we determined a genome-wide virus transcriptome and profiled their differential expression in a tissue-, virus-, and temporal class-dependent manners. Further studies identified several putative ORFs that encode hypothetical proteins containing viral mimicking conserved domains found in host interferon (IFN) regulatory factors (IRFs) and IFN receptors. This study provides the first comprehensive genome-wide viral transcriptome profiling during infection and across multiple amphibian host tissues that will serve as instrumental reference. It also presents evidence implying that ranaviruses like FV3 have acquired previously unknown molecular mimics interfering with host IFN signaling during evolution.

## 1. INTRODUCTION

Frog virus 3 (FV3) is a large (∼105 kb), double-strand DNA (dsDNA) virus belonging to the Ranaviruses genus (family *Iridoviridae)*, which comprises a group of emerging viruses that infect cold-blooded animals including amphibians, fish and reptiles [1, 2]. FV3 infections were first reported in leopard frogs in 1960s, and several virus isolates were obtained from cultured tissues/cells of both healthy frogs and tumor-bearing ones with renal carcinoma [1–3]. This implied a tumorigenic potential; however, further studies demonstrated no etiological association of FV3 with the renal oncogenesis [1–3]. On the other hand, the association of FV3 with apparently healthy frogs indicates host-adaptive transmission and persistence, and may *de facto* cause diseases in other susceptible stages during the amphibian life cycle [4]. More and more studies have implicated Ranaviruses to the decline of amphibian populations worldwide across much of their habitat regions [5–8]. FV3 represents the most frequently reported iridovirus for anurans. In North America, FV3 is widespread in wild amphibians and the only ranavirus detected in turtles [6,9,10]. A recent study detected different FV3 lineages in wild amphibians in Canada, and these new FV3 isolates appears to have undergone genetic recombination with the common midwife toad virus (CMTV) [9]. CMTV represents another ranavirus to affect various amphibians and reptile species and cause mortality events throughout Europe and Asia [9, 10]. These findings reinforce the urgency to study ranaviral biology to face the bio-ecological threat from current catastrophic amphibian decline and negative impacts in aquaculture [1–10].

Among various ranaviruses accounting for epizootics in amphibians, fish and reptiles, FV3 is the best-characterized model and the prototype of the genus ranavirus [1, 2]. Historically, FV3 studies have provided insights into the ranavirus biology including relevant characterization of highly methylated and phage-like genetic DNA, two-stage viral genome replication, temporal transcription, and virus-mediated arrest of host response [2, 11]. Prompted by early studies of FV3’s DNA synthesis occurring at the two-stage fashion between the nucleus and the cytoplasm [11], more comparative genomic studies of large nuclear and cytoplasmic DNA viruses (NCLDVs) of eukaryotes have revealed the monophyletic origin of four viral families: poxviruses, asfarviruses, iridoviruses, and phycodnaviruses [12–14]. As recent proposals extend NCLDVs to include three other taxonomic families (*Ascoviridae*, *Marseilleviridae*, *Mimiviridae*) and new founding members of other types of giant dsDNA viruses, advances in ranavirus research contribute to delineate viral evolution and host tropism diversity among iridoviruses and NCLDVs, which can unveil evolutionary links among viruses associated with different cellular life forms [15, 16]. However, in spite of the general characterization of FV3 replication and infection, the transcriptomic profile of the many viral genes and precise roles of most viral proteins of FV3 and most other ranaviruses remain elusive.

Early pioneering studies had resolved 47 viral RNAs and 35 viral proteins using gel electrophoresis in FV3-infected fish cells, and temporally classified them into early, immediate delay, and late genes along the viral infection cycle [17, 18]. The first report of transcriptomic analysis used both microarray hybridization and RT-PCR validation to examine the expression of all 98 coding genes (or open reading frames, ORFs) as annotated in the FV3’s reference genome [19]. In that study, Majji et al. identified 33 immediate early (IE) genes, 22 delayed early (DE) genes, 36 late (L) viral genes, while seven remaining genes were undermined. As postulated for temporal class of FV3’s genes in general, early genes (including both IE and DE) encode putative regulatory factors, or proteins that act in nucleic acid metabolism and immune regulation, whereas products of L genes are involved in the virion packaging, assembly and cellular interaction for viral release [19]. Notably, all these previous FV3 gene transcription studies were performed *in vitro* using a fathead minnow (FHM) fish cell line model [17–19]. Thus, to date FV3 transcriptomic profiling *in vivo* in infected host is lacking.

To complement recent virome studies and novel ranavirus isolations, it is imperative to characterize *de novo* FV3’s transcriptome and conduct gene functional analysis using next generation sequencing (NGS)-facilitated metagenomics approaches [20, 21]. To establish a procedure for unbiased analyses of ranaviral gene expression at a genome-wide scale, we have performed a whole transcriptomic analysis (RNA-Seq) using total RNA samples containing both the viral and cellular transcripts from FV3-infected frog tissues. Two FV3 strains, a wild type (FV3-WT) and an ORF64R-knockout strain (FV3-Δ64R), were used for comparison [22, 23]. The ORF64R gene encode a putative interleukin-1 beta convertase containing a caspase activation and recruitment domain (vCARD) that is postulated to serve as an immune evasion factor as a result of FV3-Δ64R [22, 23]. It was, therefore, interesting to assess the viral gene response under stronger host immune pressure. Our analyses identified the expression of most annotated 98 ORFs and profiled their differential expression in a tissue-, virus-, and temporal class-dependent manners. Furthermore, we used a reverse-genetic approach to functionally identify viral putative ORFs that encode hypothetical proteins, particularly those containing viral mimicking domains analogical to that in host interferon (IFN) regulatory factors (IRFs) or IFN receptors, especially for the type III IFNs. As a cardinal antiviral mechanism diversified along tetrapod evolution, the IFN system comprises three types of IFNs (type I, II, and III) that are classified mainly based on their molecular signatures and type-specific cognate receptors [24–26]. IFNs induce diverse immune responses extensively characterized in antiviral responses and are involved in immunomodulatory process through signaling cascades *via* respective IFN receptors and various IRFs [24–27]. Previous studies have determined the key position of amphibians in IFN evolution [25, 26], and the alteration of viral infection on IFN response in FV3-infected frogs [22, 23]. The functional analyses may provide mechanistic explanation about the viral interference of IFN responses in FV3-infected cells/tissues, and provide new insights into evolutionary arm race between the ranavirus and quickly evolving amphibian IFN system [22–26]. Our study thus provides the first virus-targeted genome-wide transcriptome profiling of a large DNA virus during real amphibian host infection and uncover potential function of hypothetical proteins in the context of virus-host interaction.

## 2. MATERIALS AND METHODS

### Animals and virus

Outbred specific-pathogen-free adult (1-2 years old) frogs were obtained from the *X. laevis* research resource for immunology at the University of Rochester (http://www.urmc.rochester.edu/mbi/resources/xenopus-laevis/). All animals were handled under strict laboratory and University Committee on Animal Resources regulations (approval number 100577/2003-151). Two FV3 strains, a wild type (FV3-WT) and an ORF64R-desrupted strain (FV3-Δ64R), were used for comparison. Frog virus 3 (FV3) stock preparation and animal infection were conducted as previously described [22, 23]. In brief, fathead minnow (FHM) cells (ATCC^®^ CCL-42) or baby hamster kidney-21 (BHK-21) cells (ATCC^®^ CCL-10) were maintained in suggested medium (DMEM or MEM; Invitrogen) supplemented with 10% fetal bovine serum (Invitrogen), penicillin (100 U/ml), and streptomycin (100 μg/ml) at 30°C with 5% CO_2_. FV3 was grown by a single passage in FMH or BHK-21 cells and virus stocks were purified by ultracentrifugation on a 30% sucrose gradient and the virus load was assessed by plaque assays on BHK-21 monolayer under an overlay of 1% methylcellulose (ATCC^®^ CCL-102). Virus stocks were titrated using plaque assays of serially diluted viral stocks on BHK-21 monolayers to express as plaque forming units (PFU) as previously described [22, 23].

### Animal infection and tissue sampling

Adult frogs at comparable age and body-weight were randomly allotted into mock controls and infected groups (n = 5/group). Animal infections were conducted by intraperitoneal (i.p.) injection of each animal with FV3-WT or FV3-Δ64R at 1 × 10^6^ PFU in 100-μl amphibian phosphate-buffered saline solution (APBS) or only APBS for mock controls. At 0, 1, 3, and 6 days post-infection, animals were euthanized by immersion in bicarbonate buffered 0.5% tricaine methane sulfonate (MS-222), and indicated tissues were sampled and pairwise allotted for classical viral titration and gene expression analyses, and the samples of 3 day post infection were selected for further unbiased or targeted transcriptomic studies as diagramed in Figure 1 [22, 23].

**Figure 1.**
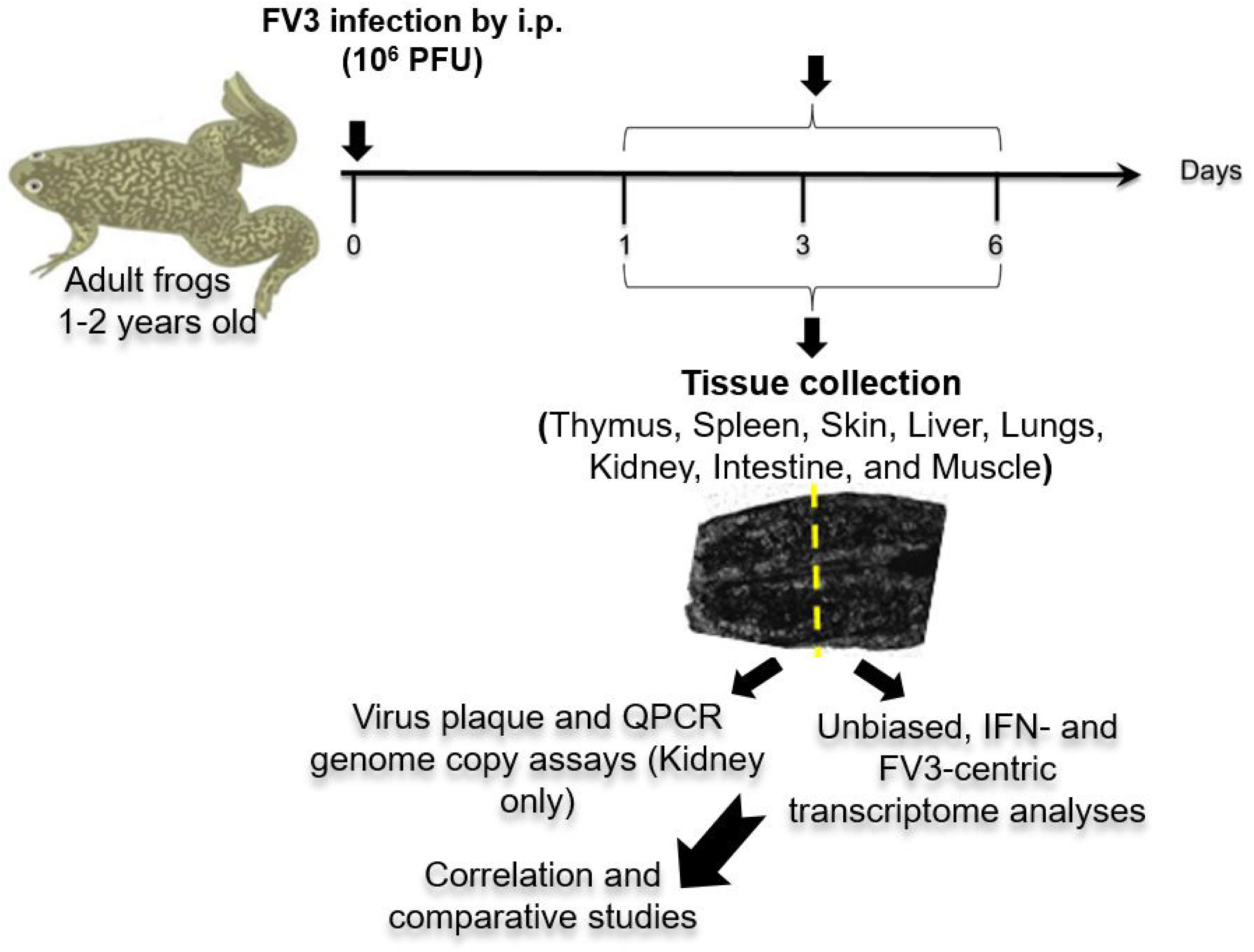
Experimental sample collection and processing. Adult healthy frogs (*Xenopus laevis,* n= 5/group) at 1-2 years old were infected intraperitoneally (i.p.) with a wild-type (WT) Frog Virus 3 (FV3) or an Orf64R-disrupted laboratory strain (Δ64R) at the dose of 10^6^ plaque forming unit (PFU). Indicated tissues were sampled and pairwise allotted for classical viral titration and gene expression analyses, and the samples of 3 day post infection were selected for further unbiased or targeted transcriptomic studies as illustrated and described in the text.

### DNA/RNA extraction and qPCR FV3 gene copy assays

Total RNA and DNA were extracted from frog tissues and cells using a TRIzol reagent (Invitrogen) for PCR-based assays or a column-based RNA/DNA/protein purification kit (Norgen Biotek, Ontario, Canada) for transcriptomic analysis. RNA integrity and concentration were evaluated with a NanoDrop 8000 spectrometer (NanoDrop, Wilmington, DE) and an Agilent 2100 Bioanalyzer (Agilent Technologies, Santa Clara, CA) to ensure RNA samples with A260/A280>1.8 and RNA integrity number (RIN) >7.0 qualified for construction of sequencing libraries [28, 29].

Quantitative PCR (qPCR) analysis was performed using 150 ng/reaction of DNA templates in an ABI 7300 real-time PCR system and PerfeCta SYBR green FastMix, ROX (Quanta). To measure FV3 genome copy number based on detection of FV3gorf60R, which encodes a viral DNA polymerase II (Pol II), a qPCR was performed against a standard curve generated using a serially diluted template DNA containing 10^1^ to 10^10^ vDNA Pol II DNA copies cloned in a plasmid as previously described [22, 28].

### Transcriptomic assays (RNA-Seq)

RNA samples used for RNA-Seq sequencing library preparation were pooled from three qualified extractions of each group as indicated above. For sequencing libraries construction, mRNA purification, fragmentation, construction of sequencing libraries and sequencing were performed using the Illumina Pipeline (Novogene, Sacramento, CA). To ensure representation of ranaviral transcripts potentially lacking poly-adenylated tails, we used a kit for isolation of total RNA (Catalog # 20040529, Illumina). Approximately 40 M clean reads per sample were generated for genome-wide transcriptomic analyses. The trimmed reads were further assembled and mapped to the Reference genome/transcripts of *X. laevis* or FV3 virus through Xenbase (http://ftp.xenbase.org/) or NCBI genome ports (ftp://ftp.ncbi.nlm.nih.gov/ genomes/all/GCF), respectively. Only data for the virus-targeted transcriptome was reported here. The workflow of RNA-Seq analysis and data representative of general quality and comparability of the transcriptome data are shown in Supplemental Figure 2. Software used for reads mapping, quantification, differential analysis, and sequential gene ontology (GO) and pathway analysis, was listed in Table 1. Significantly and differentially expressed genes (DEGs) between two treatments were determined using DeSeq and edgeR packages and visualized using bar charts (FPKM) or heatmaps (Log2 fold ratio) as previously described [29]. The transcriptomic dataset was deposited in the NIH Short Read Archive linked to a BioProject with an accession number of PRJNA705195.

**Table 1.**
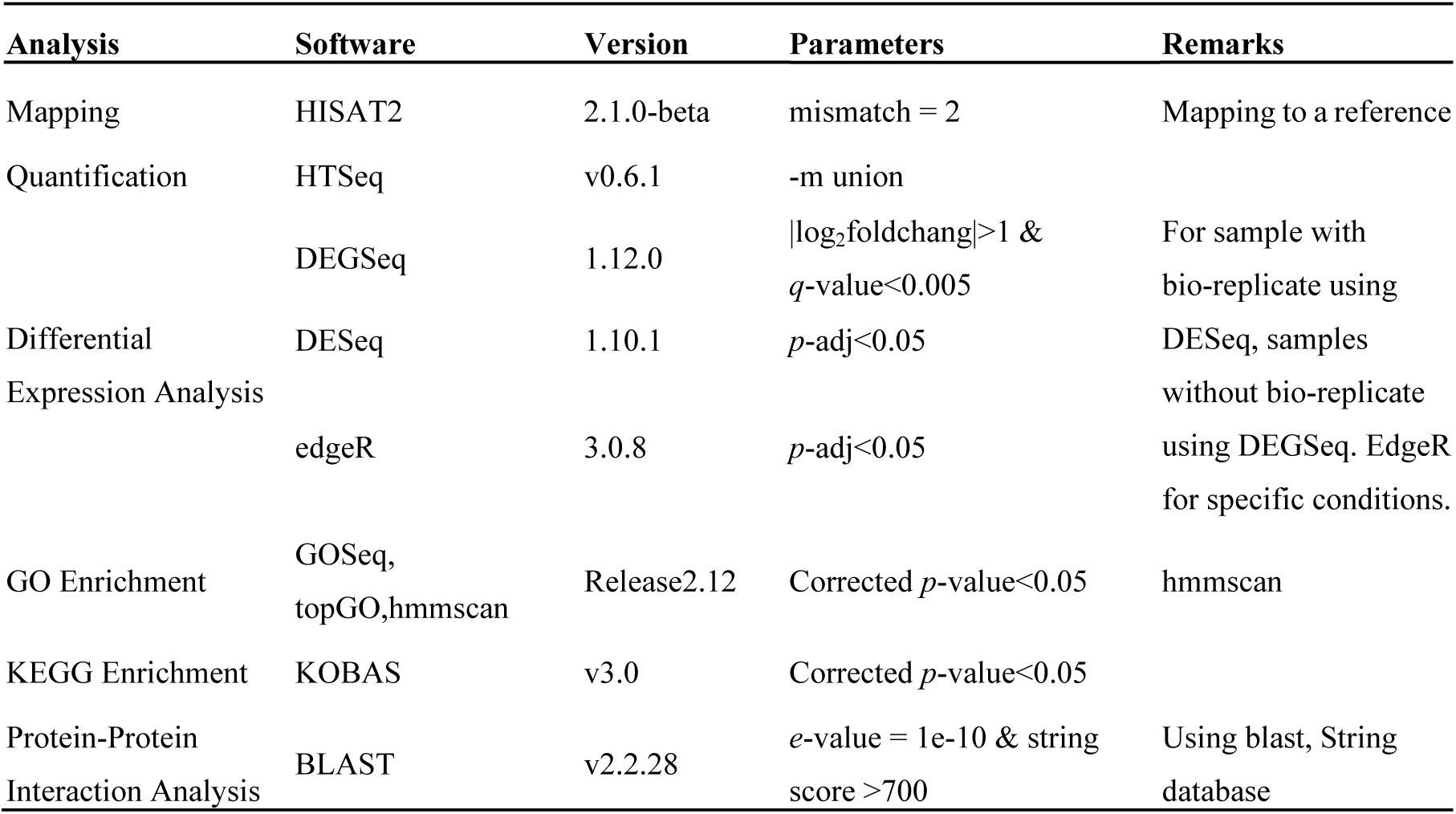
Software List for data bioinformatic analysis.

### Novel viral gene prediction and functional analysis

We conducted extensive sequence- or pattern-based Blast searches against FV3 reference genome (GenBank Accession No. NC_005946.1) using conservative domains in *Xenopus* proteins of IFN signaling, especially those of IFN receptors and IFN regulatory factors (IRFs). The Blast searching programs were mainly through NCBI Blast portal at https://blast.ncbi.nlm.nih.gov/Blast.cgi with Expect threshold (E-value) adjusted to 1. Only viral proteins, which showed E-value less than 0.5 and contain regions spanning most or full coverage of the functional domains in aligned host proteins were selected for further simulation analyses. Further viral coding gene prediction integrated to use both programs fgenesV0 and fgenesV through http://www.softberry.com/, and annotated as novel open reading frames (Norf) if they were not annotated along the FV3 reference genome. The protein domain analysis were queried and extracted using NCBI CDD database. The full-length sequences of the predicted hypothetical ORF/proteins are provided in the Supplemental Excel Sheet. The GenBank Accession Numbers of all aligned gene/protein sequences are listed in Table 2 [25].

**Table 2.**
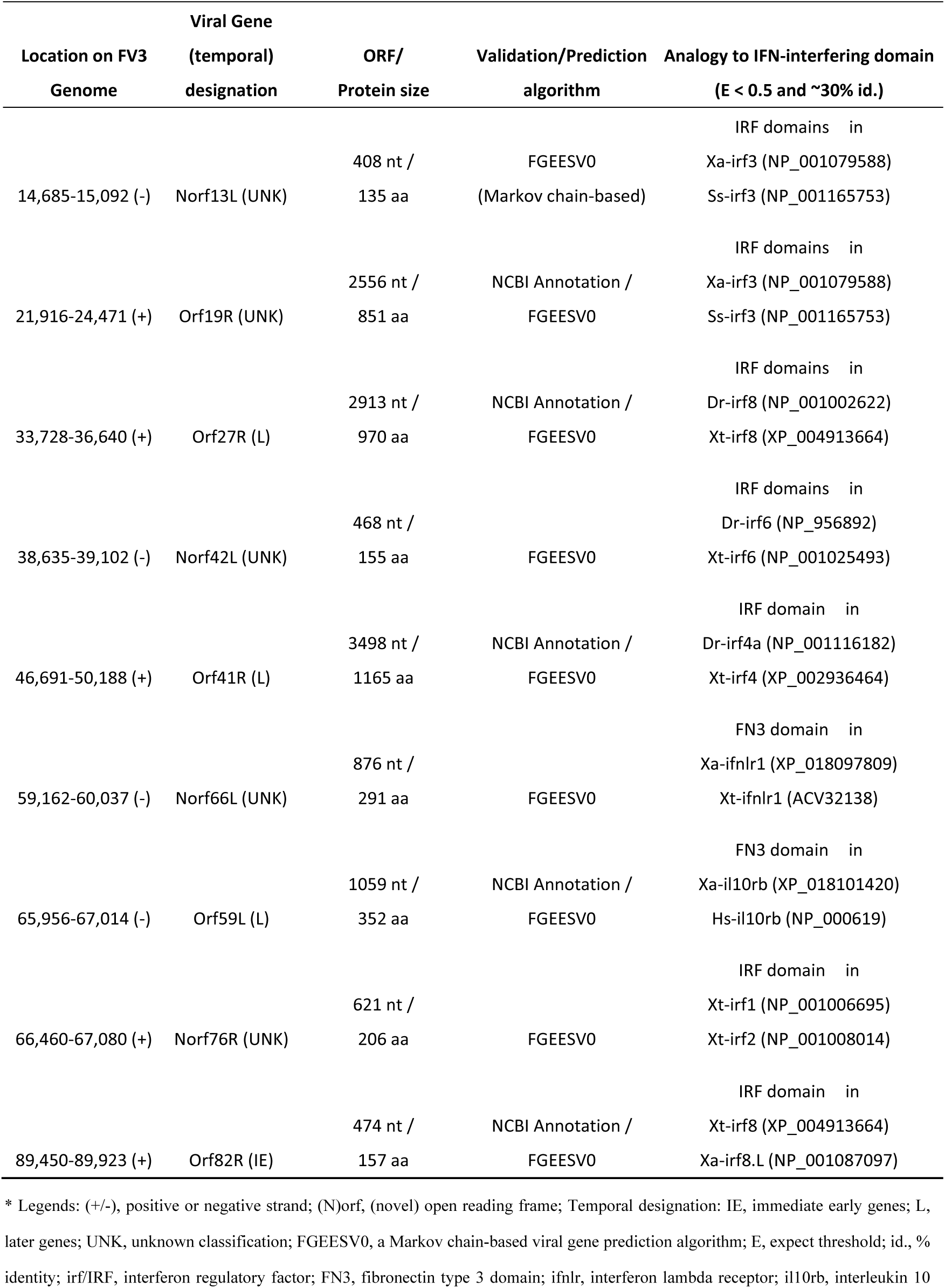
Viral genes that encode hypothetical proteins containing conserved domains potentially able to interfere with host interferon signaling*.

Multiple sequence alignments and views were done with a Jalview program. Protein structure models were simulated and visualized through combinative uses of the programs of PyMol, Chimera, and Phyre2 as described [30], and primarily a HDOCK server (http://hdock.phys.hust.edu.cn/) for protein-protein or protein-DNA docking based on a hybrid algorithm of *ab initio* free docking. Without further indication in the legends, all programs were used under a defaulted condition.

### Statistical analysis

Statistical analysis was conducted using one-way analysis of variance (ANOVA) and Tukeyʹs post hoc test. A two-sample F test was used for significant evaluation between samples/treatments. A probability level of *p*<0.05 was considered significant [25, 28].

## 3. RESULTS AND DISCUSSIONS

### FV3 infection and comparative viral determination between FV3-WT and FV3-Δ64R strains in the kidney of adult frogs

FV3 infects anuran amphibians at various developmental stages and are highly lethal in tadpoles. Adult frogs, by contrast, are more resistant to viral infection, and after viral clearance low level of quiescent viruses were isolated from apparently healthy frogs [22,23,28]. This indicates that adult frogs are more adaptive to the virus-caused deathliness [23]; and meanwhile serve as active carriers or reservoirs for the virus transmission and provide a valuable model for studying the virus-host coevolution [27]. In this context, we infected in laboratory-controlled conditions in 1-2 years old *X. laevis* frogs for well-controlled sample collection. As shown in Figure 1 and Figure 3, randomly allotted frogs were infected with either a FV3-WT or FV3-Δ64R strain. Time point were chosen based on previous published study to include an early innate immune response (1 dpi) and intermediate (3 dpi) and the peak of adaptive T cell immune response (6 dpi) [3, 4]. Intraperitoneal inoculation was used to minimize variation of initial infection and to be able to rely on previous characterization of wild type and KO FV3 infection in X. laevis [3, 4]. Both FV3 strains caused early productive infections as shown with successful viral isolation from the kidney tissue homogenates of infected frogs (Figure 3A). The rationale to use the FV3-Δ64R mutant virus deficient for a putative immune evasion gene was to evaluate to global viral gene expression response under stronger host immune pressure. As anticipated, FV3-WT was more efficient in producing infectious virions compared to the)), showing 100-1000 magnitude difference compared using a logarithmic scale of PFU at 1-6 dpi (Figure 3B). Similar infection patterns were also observed by measuring the virus genome copy number based on quantitative detection of the viral Orf60R gene, which encodes a virus DNA polymerase II unit (Pol II) (Figure 3C). However, the viral genome copy number of FV3-WT kept increasing through 1-6 dpi, whereas, FV3-Δ64R genome copies reached a higher level than the wild type at 1 dpi and keep at a similar level through the tested period without much increase (Figure 3C). From daily observation throughout the infection process, infected frogs behaved similar to the mock-infected controls and no outliers per clinical observation or virus diagnosis were identified, which were averagely qualified for sample collection as designed for further transcriptomic process (Figure 1 and 2).

**Figure 2.**
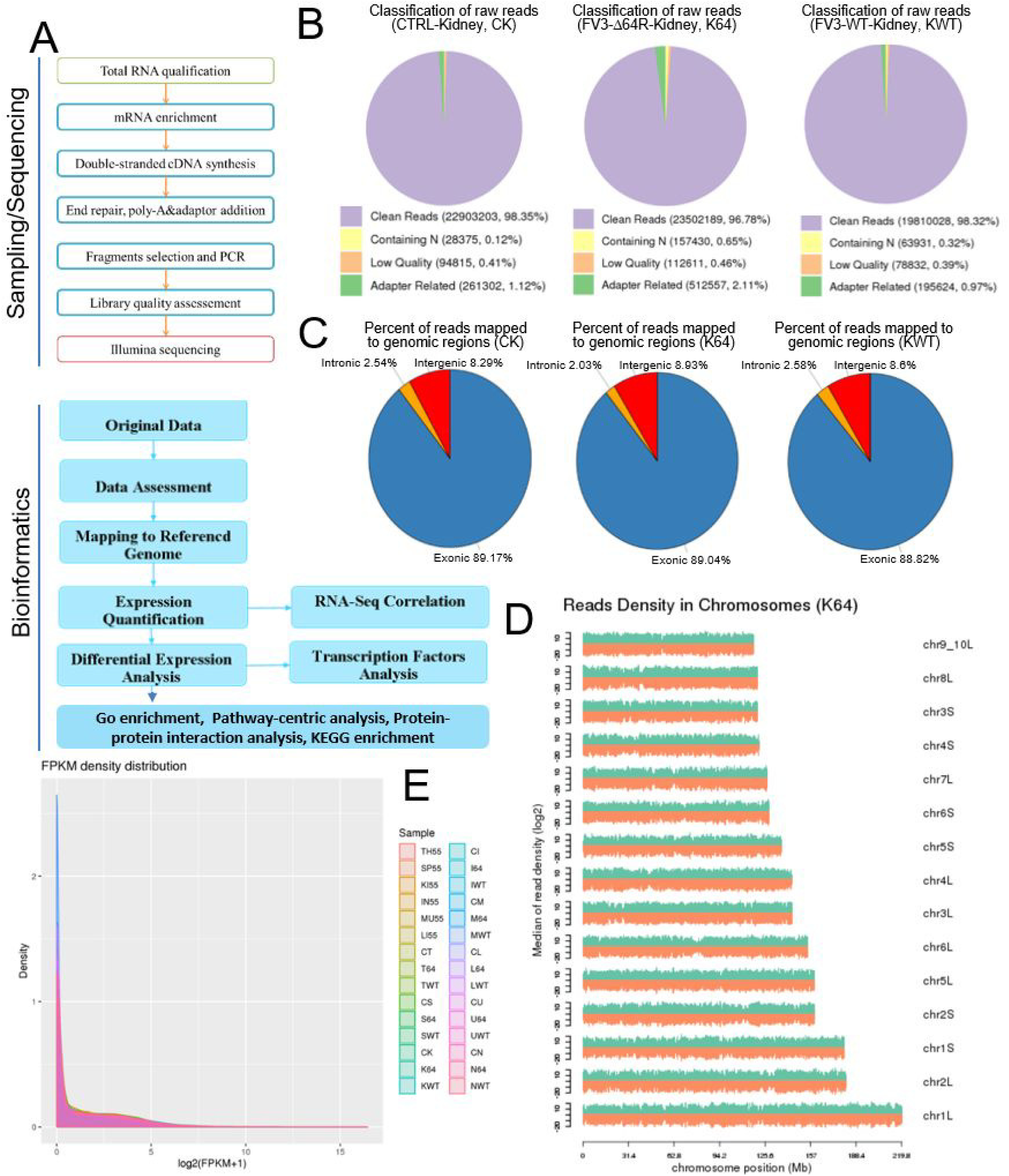
The workflow and data showing general quality and comparability of the RNA-Seq transcriptome data. (A) Workflow of the Illumina RNA-Seq procedure and bioinformatic analysis. (B), (C) and (D) The composition and comparability of raw reads (B), percent of reads mapped to genomic regions (C), and reads density and coverage in Chromosome (D) in representative samples. Very similar reads coverage and distribution were comparatively obtained in all tissue samples in general. (E) FPKM density distribution to show different gene expression levels under different experiment conditions. FPKM distribution, the x-axis shows the log_10_(FPKM+1) and the y-axis shows gene density. FPKM, Fragments Per Kilobase Million for paired-end RNA-Seq. Please refer the determination for sample types of abbreviations in the text and figure legends as indicated.

**Figure 3.**
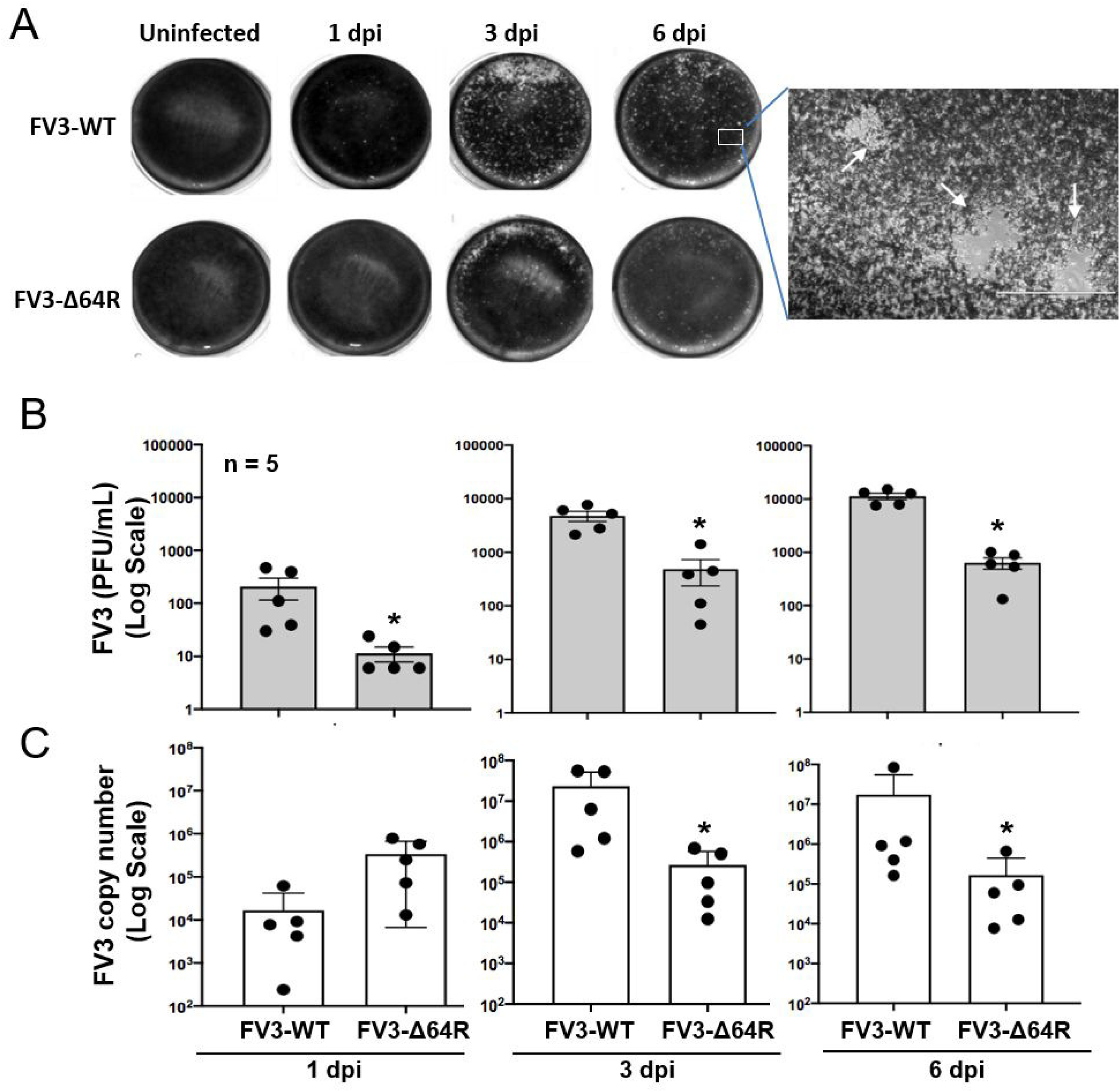
Viral plaque assays and genome copy number detection by quantitative PCR (QPCR). (A) For plaque assays, 1 ml of the virus-containing supernatant of each individual kidney homogenate was used to inoculate A6 cells (ATCC® CCL-102™) and FV3 plaques were counted at 1, 3, and 6 day post infection (dpi) and imaged for representative wells. (B) Virus titers were calculated as average PFU/ml for each group/treatment. (C) Virus genome copies were determined by QPCR in comparison to titrated copies of clones FV3gorf60R fragment in 150 ng DNA per reaction as described. *p < 0.05, n = 5 for (B) and (C).

### Virus-targeting transcriptome analysis and difference dependent on tissue types and FV3 strains

FV3’s genome encodes 98 putative coding genes (FV3gorf1-98) as annotated along its reference genome [19]. Previous microarray analysis plus RT-PCR validation determined the expression of all the 98 FV3 ORFs, indicating a full-genome transcribing capacity during FV3 infection in the FHM cell line [19]. Consistent with the microarray analysis, our comprehensive unbiased transcriptome analysis based on *de novo* deep sequencing revealed FV3 gene-specific reads spanning the full FV3-genome at ∼10X depth in samples from the infected intestine (FV3-WT only), liver, spleen (FV3-Δ64R only), lung and particularly kidney (Figure 4). In addition, partial genome coverage or regional detection were obtained in FV3-infected muscle, skin and thymus tissues (Figure 4). Importantly, no FV3-specific reads were detected from all sham-infected control tissues, ruling out cross-contamination and validating our sample handling procedures.

**Figure 4.**
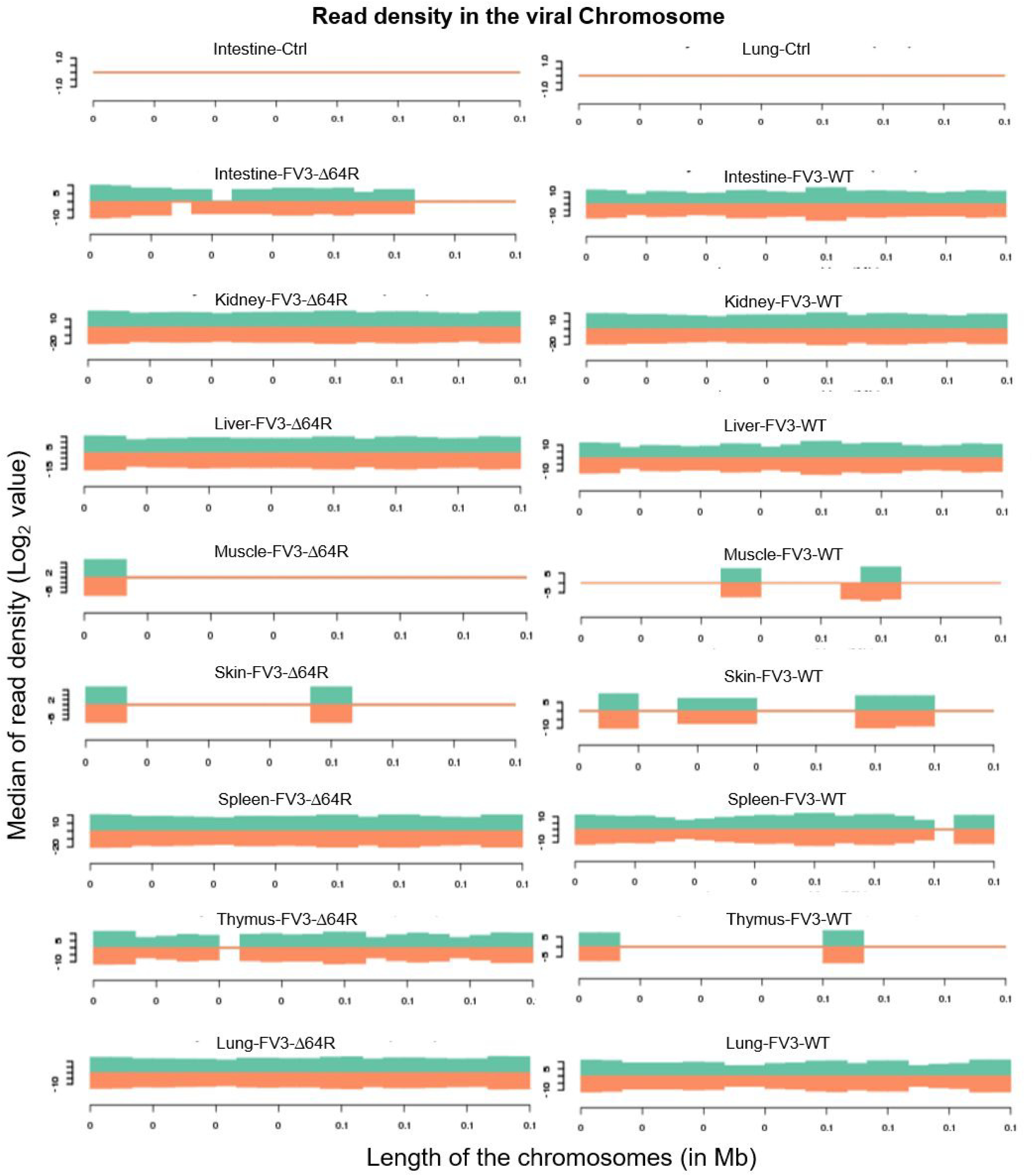
Virus-targeted transcriptome analysis. Shown is distribution plots of mapped reads in FV3 genome (GenBank Accession No. NC_005946.1). The X-axis shows the length of the genome (in Mb, 0.105 Mb of FV3), and the Y-axis indicates the log_2_ of the median of read density. Green and red indicate the positive and negative strands, respectively. Note, no FV3 transcript read were obtained from all control (Ctrl) non-infected samples (shown only those from the intestine and lung).

Statistical analyses of the RNA-Seq results revealed several interesting aspects: (1) FV3 may maintain a more complex transcript mixture *in vivo* than in a uniform cell line related to unsynchronized infection stages upon diverse cell types in tissues. Given that the viral transcripts were significantly detected in the kidney, spleen, liver, lung, thymus and intestine, we conclude that the differential viral gene expression in these tissues resulted primarily from systemic virus-host interaction initiated from intraperitoneal injection. (2) FV3-specific reads were significantly enriched in an increasing order in the intestine, lung, liver, spleen and predominantly in kidneys, but much less abundant in the skin and muscle where FV3 replication may be negligible. (3) The transcript profiles of FV3-Δ64R mutant were nearly identical to the FV3-WT in the kidney, but qualitatively and quantitatively very different in other tissues. (4) The unbiased transcriptome study detected transcripts of FV3 genome almost equivalently along both positive and negative strands, which confirms the existence of viral coding genes at both strand orientations (Figure 4). Further co-expression-venn analysis confirmed a virus-strain (FV3)-dependent difference of gene transcription among most tested tissues, as both WT and FV3-Δ64R shared near identical transcript profiles of all annotated 98 FV3gorfs in the kidney. Interestingly, although WT FV3 genes were more efficiently transcribed than FV3-Δ64R in the intestine, skin and kidney, the recombinant mutant virus actually had much more transcripts in the thymus, liver, lung and particularly spleen. This implies that the disruption of the FV3gorf64R gene, which encodes a putative interleukin-1 beta convertase containing caspase recruitment domain (vCARD), may change viral transcription dynamics and the tissue/cell tropism of FV3 infection in amphibians (Figure 4 and data not shown). As shown in Figure 5, the Pearson correlation analysis demonstrated a general low cross-sample correlation, except that between the FV3-Δ64R-infected spleen and kidney, further indicating both tissue types had a top priority to support FV3 infection and full-scale of gene expression (Figure 5).

**Figure 5.**
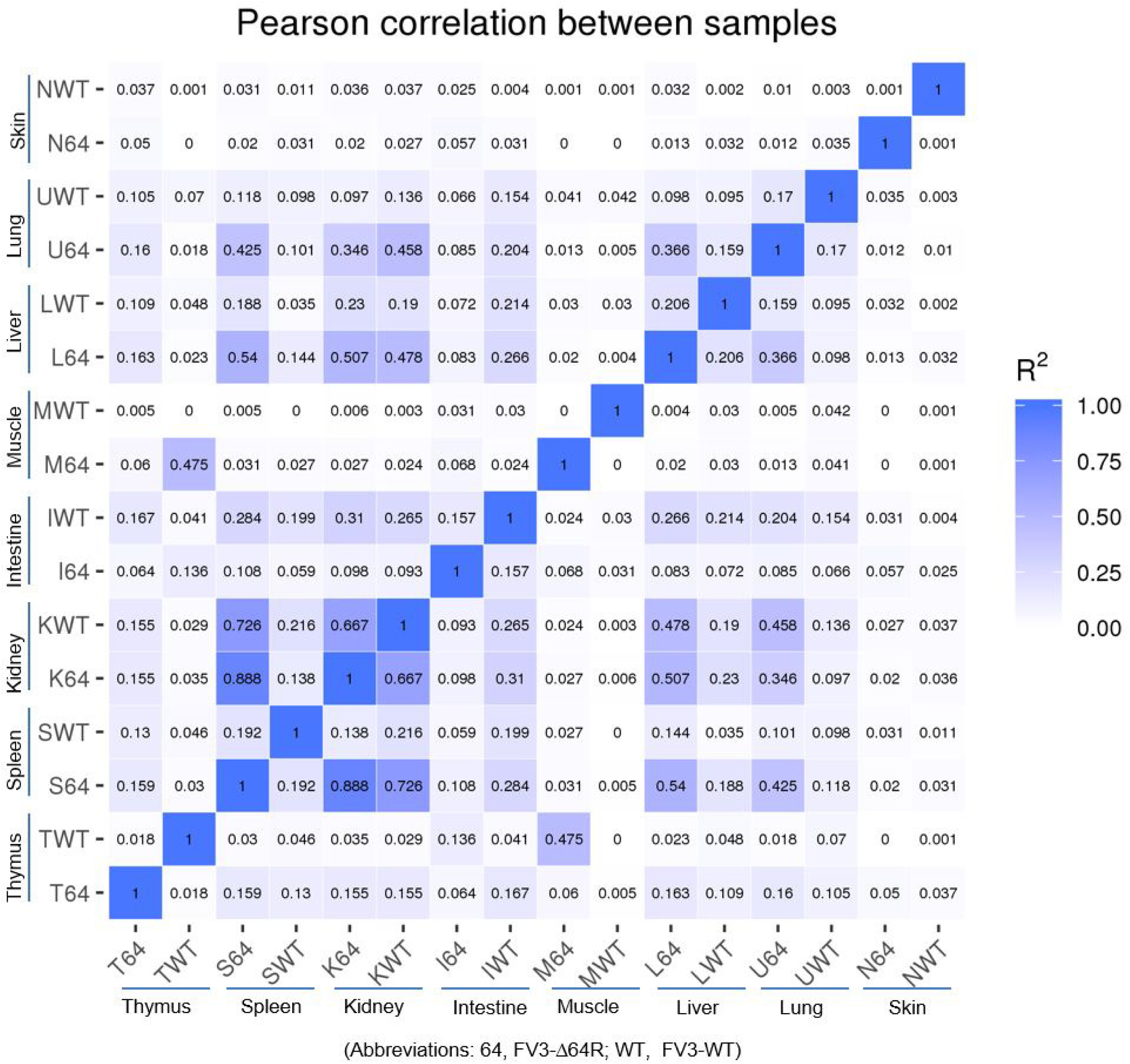
RNA-Seq correlation among the samples. Heat maps of the correlation coefficient between samples are shown. Numbers indicate the square of the Pearson coefficient (R^2^). The closer the correlation coefficient is to 1, the greater the similarity of the samples. Generally low R^2^ values indicate dramatic difference between different tissues and two FV3 strains.

### Genome-wide differential expression analysis of FV3 coding genes

Figure 6 presents an heatmap and cluster analysis of differential expression based on all 98 putative ORFs of the annotated FV3 genome. Using the FPKM (fragments per kilobase of transcript per million mapped reads) values as standardized for paired-end RNA-Seq analysis, results showed a tissue-dependent expression of various viral genes across the genome-wide ORF panel. Again, most FV3 genes were highly expressed in kidney and spleen, but quite few genes also showed high expression levels in other tissues including intestine, liver, thymus and lung (Redline framed in Figure 6). Within major gene clusters with similar tissue expression patterns, the correlation to their functional relevance or temporal class was observed to some extent. However, this should not be considered as a general reference for interpreting viral gene function within each cluster. Because even when genes shared cross-tissue expression patterns, they apparently had different putative functions and were ascribed into different temporal classes (Figure 6). Again, most viral genes had differential expression patterns between the FV3-WT and FV3-Δ64R strains, suggesting that the putative cCARD gene play an important role in mediating virus-host interaction, which influence viral transcription. Among 98 annotated FV3gorfs, no specific reads mapped to three viral genes of FV3gorf-55L, -30R and -68R (black rows in Figure 6), which implies that during infection *in vivo* these ORFs were not transcribed or underwent transcript decay in the tested tissues. However, this does not exclude their potential expression in other tissues/cell types and/or other host species since they have been previously detected in the FHM cell culture by microarray [19]. This underscores the need of extensive comparative analysis of the virus transcriptome at different time points post the viral infection in both tadpoles and adult frogs of different species using the established NGS platform (Figure 2 and Table 1).

**Figure 6.**
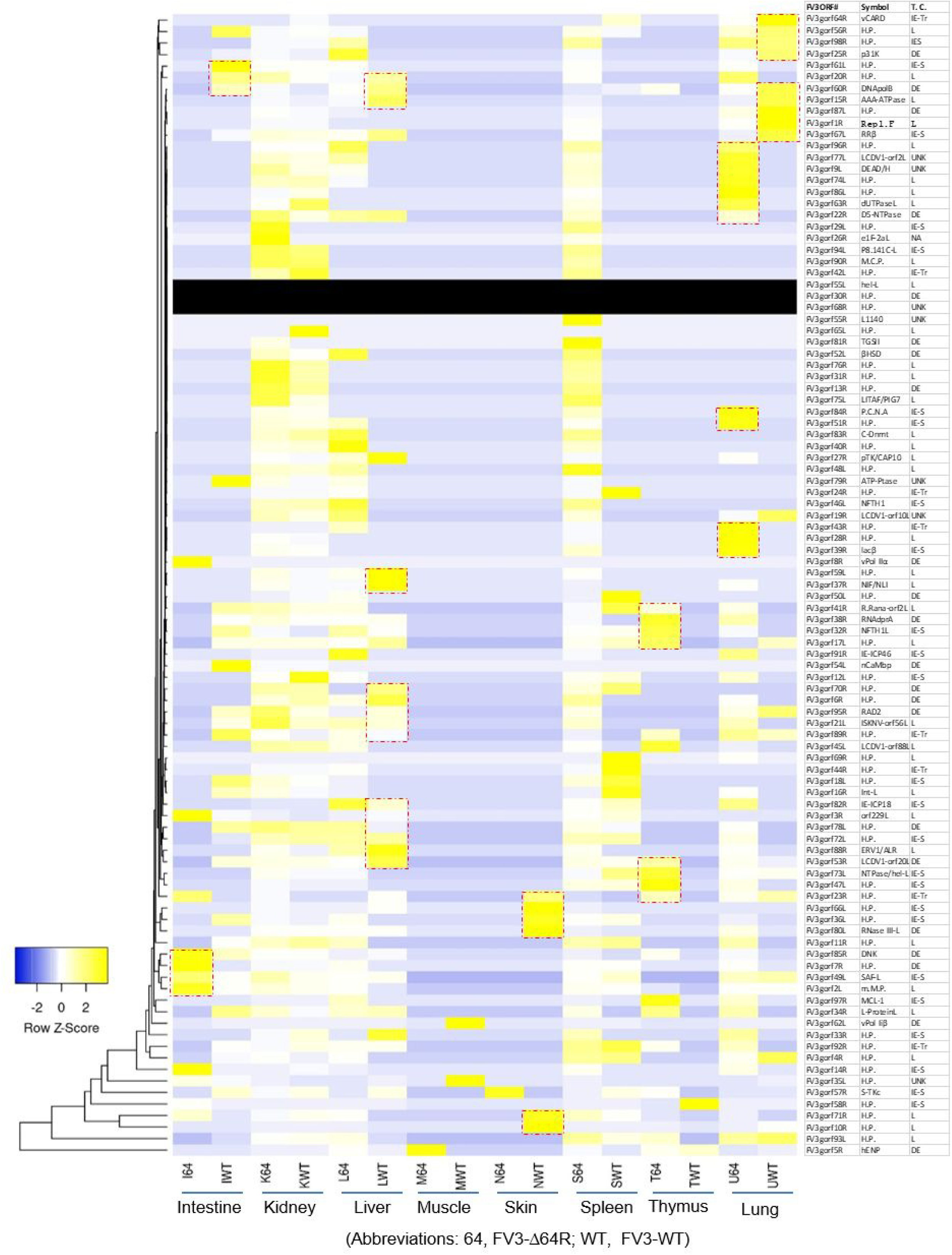
Heatmap and cluster analysis of differential expression of all validated/putative ORFs for coding genes along FV3 genome. FPKM values were used as for paired-end RNA-Seq per differential gene expression and cluster analysis. Clustered using the log_10_(FPKM+1) values. Yellow denotes genes with high expression levels, and blue denotes genes with low expression levels. The color ranges from yellow to blue represents the log_10_(FPKM+1) values from large to small. The ORFs sit in a same or close clusters have similar expression patterns across the samples. The black rows indicate that the expression of the corresponding ORFs has not been detected, implying a silent or nonproductive transcription. The table at the right lists the 98 annotated open reading frames (ORFs) in FV3 genome, and corresponding designation of gene symbols as defined under the GenBank FV3 reference genome (NC_005946.1). Redline framed indicate higher expressed gene clusters in the infected tissues other than the kidney and spleen, where FV3 primarily showed high expression in general. Other abbreviations: FPKM, fragments per kilobase of transcript per million mapped reads; H.P., hypothetical proteins; T. C., temporal class.

### Differential expression analyses according to the temporal class of FV3 coding genes

As mentioned, it is widely assumed that early (E) genes include those encoding regulatory factors or proteins that mediate nucleic acid metabolism and immune interaction, whereas late (L) genes primarily take part in DNA packaging, and virion assembly and releasing. Transcripts of FV3’s E genes are further classified into immediate early stable messages (IE-S), immediate early transient messages (IE-Tr), and delayed early (DE) transcripts [17–19]. Figure 7 categorized the differential expression of FV3 ORF coding genes based on their temporal classification as determined above. Similar tissue- and virus-strain dependent expression patterns were clearly demonstrated with 100-1000 fold higher viral transcripts of most temporal classes in the kidney (both strains), spleen (FV3-Δ64R only), and DE in the liver (FV3-Δ64R only). The temporal class exhibiting the most differential expression pattern was the DE genes. Notably, DE transcripts were: (a) dramatically different in kidneys between FV3-Δ64R (comparably as high as the overall average) and FV3-WT (significantly lower than the overall average); (b) significantly higher on average than the other temporal classes in the infected livers; (c) significantly lower than other temporal classes in the FV3-Δ64R-infected spleen and lung, but higher in the thymus (Figure 7). These observations indicate that FV3-WT has an optimal tissue-specific regulation (or inter-tissue collaboration) of DE transcription between the kidney and livers, whereas FV3-Δ64R seems deficient in this capacity, suggesting that the FV3gorf64R gene play a critical role in regulation of DE gene expression in FV3-infected frogs.

**Figure 7.**
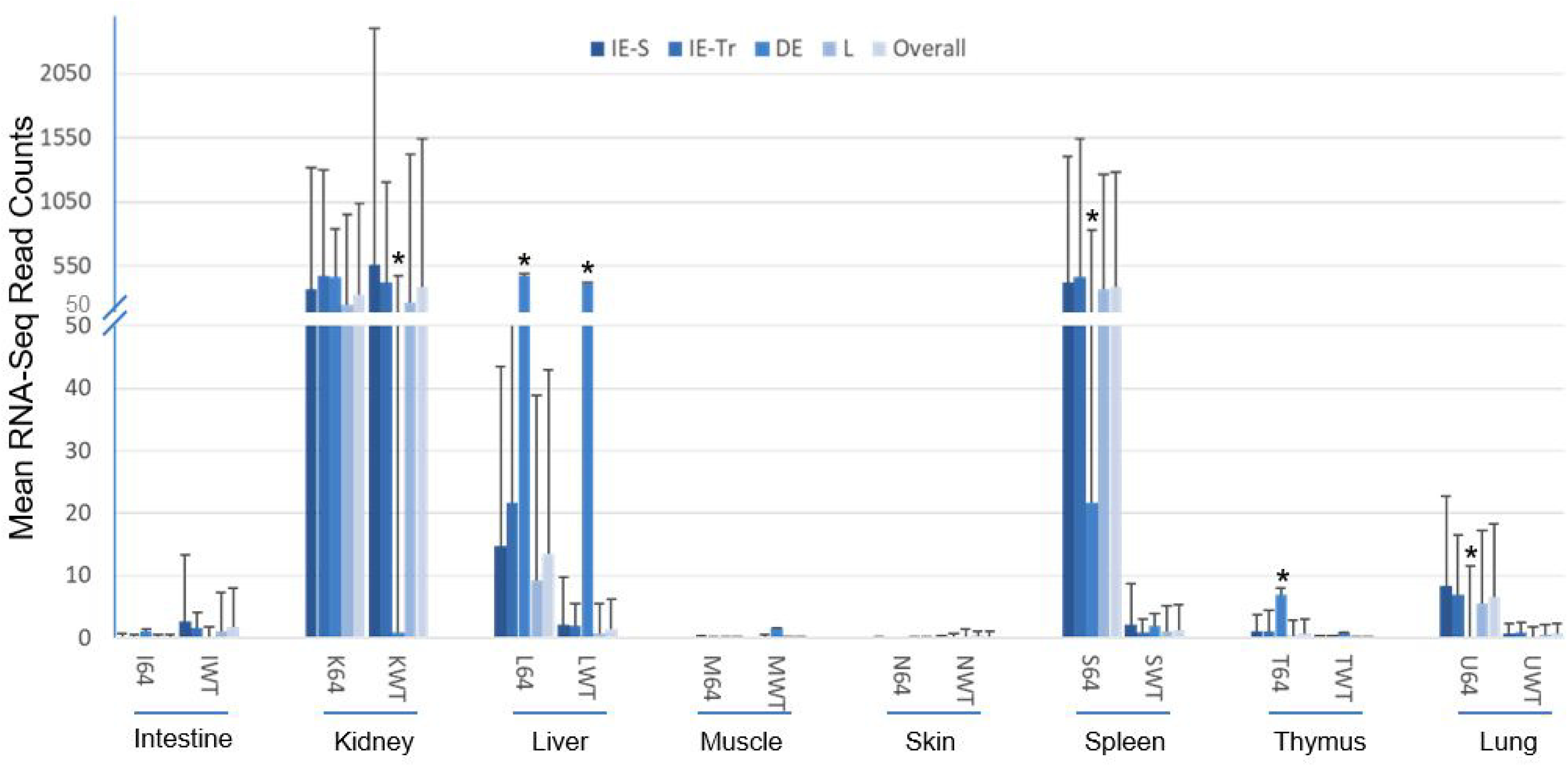
Differential expression of FV3 ORF coding genes based on their temporal classification in the viral infection. FV3 gene expression is temporally regulated in a coordinated fashion leading to the sequential appearance of immediate early (IE), delayed early (DE), and late (L) viral transcripts. The IE genes include immediate early stable messages (IE-S) and immediate early transient messages (IE-Tr). Transcriptomic analysis of the expression of the total 98 annotated FV3 ORF coding genes (*Xenbase*) indicates a differential expression pattern dependent on the gene class, virus strain, and especially tissue types. *, p<0.05, n>10, compared to the overall average. Other abbreviations: 64, FV3-Δ64R; WT, FV3-WT; Overall, genes in all the classes.

Figure 8 shows differential expression profiles of individual genes between FV3-Δ64R and FV3-WT infected groups. Although many genes had comparable expression levels between the FV3-Δ64R and FV3-WT, the average transcript counts of the WT strain were lower, particularly of the DE and L classes. Because mutant virus-infected tissues contained less infectious virions (Figure 1), it is plausible that FV3-Δ64R underwent less efficient virus packaging and resulted in higher accumulation of gene transcripts [19, 23]. In Figure 9, we present the averaged expression levels of all annotated FV3 genes sorted by decreasing order across and within each temporal gene classes. Data show that E gene group had a two-fold higher group-wide median value than L genes. It is noteworthy that the genes with an expression level close to the group means (framed by the blue dash line), are likely to serve as better gene markers for estimation of viral genome copies by classical QPCR [22,23,38].

**Figure 8.**
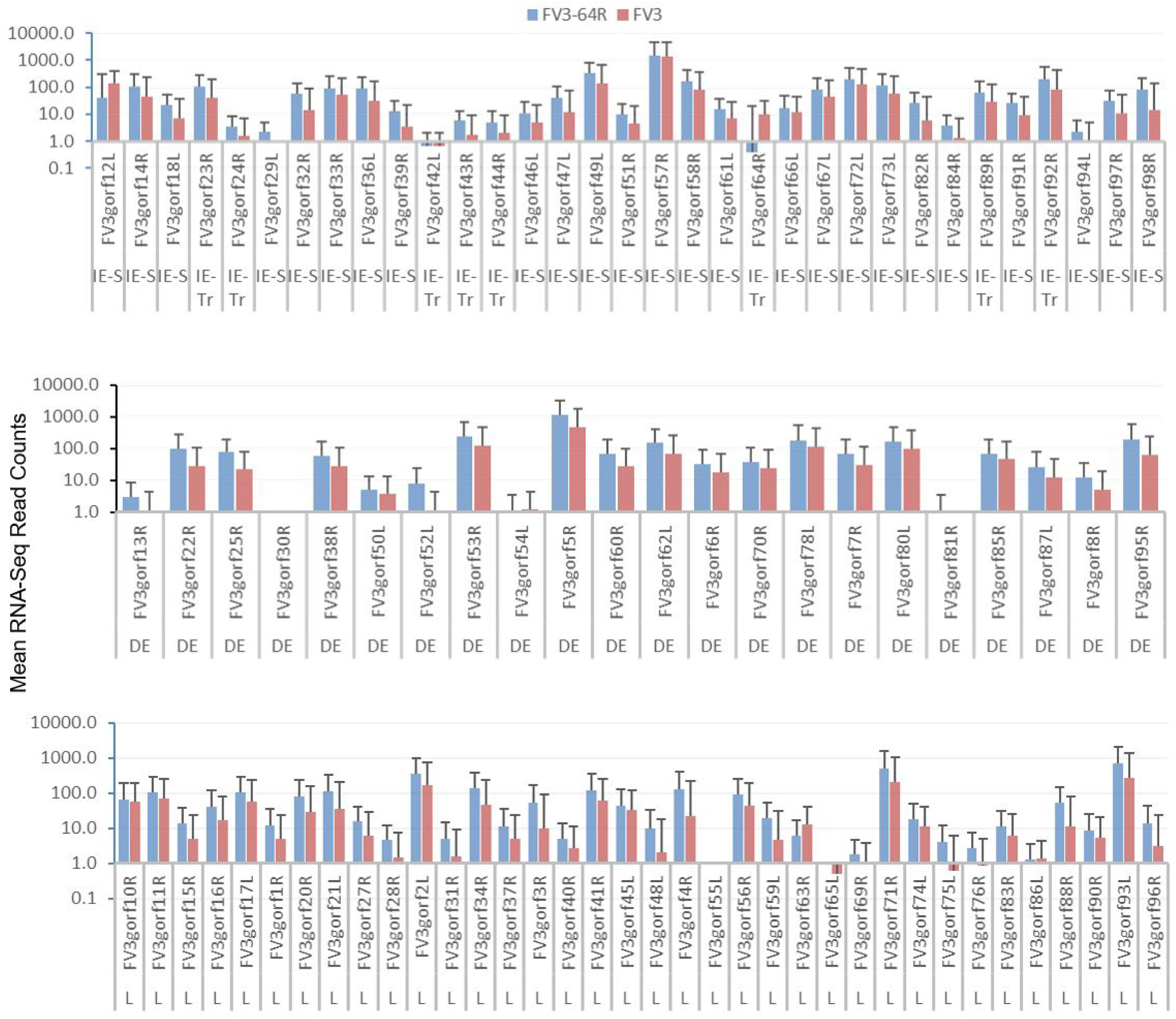
Differential expression of individual genes of FV3 between the FV3-Δ64R and FV3-WT infected groups. The genes are categorized based on their sequential classes as immediate early (IE), delayed early (DE), and late (L) viral transcripts. RNA-Seq reads are presented as averages across the eight types of different tissues to demonstrate the differential expression in dependent on the gene and virus strain. Note that the Y-axis is in a log_10_ scale. Other abbreviations: 64, FV3-Δ64R; IE-S, immediate early stable messages; IE-Tr, immediate early transient messages; WT, FV3-WT.

**Figure 9.**
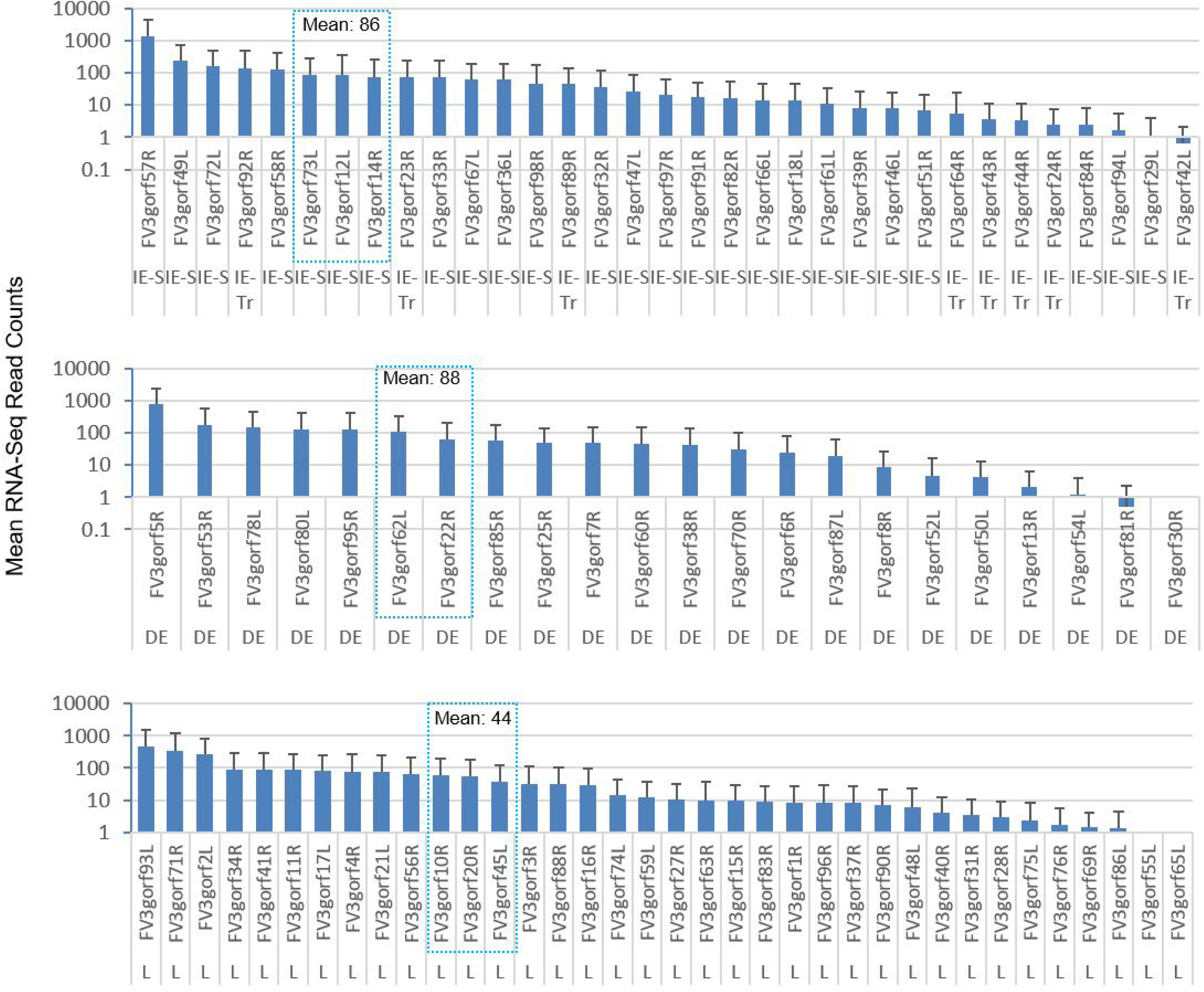
Sorted expression levels of individual FV3 ORF coding genes to show the relative expression order across and within each temporal gene classes. The genes are categorized based on their sequential classes as immediate early (IE), delayed early (DE), and late (L) viral transcripts. RNA-Seq reads are presented as averages across all samples to demonstrate the differential expression based on each gene in general at 3 day post infection. Note that the Y-axis is in a log10 scale, and the difference of the mean values between each temporal gene classes. The genes, which have an expression level close to the group means (framed by the blue dash line), should serve better gene markers for estimation of viral genome copies for classical QPCR detection. Other abbreviations: 64, FV3-Δ64R; IE-S, immediate early stable messages; IE-Tr, immediate early transient messages; WT, FV3-WT.

### Detection of interferon regulatory factor domain or Fibronectin type 3 domain in several FV3 hypothetical proteins

There are several ways to functionally characterize viral coding genes: 1) gene expression analyses; 2) inferred gene function from phylogenic, bioinformatics and/or biochemical studies; 3) determining gene involvement in viral pathogenesis or host response using such as genetic manipulation by loss-of-function or gain-of-function studies; and 4) targeted manipulation to regulate viral pathogenesis and host immunity for vaccine design. While FV3 is one of the best-studied ranaviruses, the function of most viral genes still remains poorly investigated. This is particularly the case for viral interference with the host antiviral IFN system [4,22,23]. Therefore, this study was designed to conduct for characterization of FV3 coding genes at the both the first and second aspects, i.e., establishing a high-throughput system for gene expression analyses and functional prediction for examining viral antagonistic mechanism against amphibian IFN system.

Several FV3 genes were previously assessed using FV3 knockout mutagenesis including FV3−Δ64R, FV3−Δ52L defective for putative vCARD-like protein and a β-hydroxysteroid dehydrogenase (βHSD) gene homologs respectively as well as FV3−Δ82R with a disrupted gene encoding an IE-18kDa protein (ICP18), [22, 23]. Examination of viral loads and responsive cytokine gene expression in macrophages and kidney tissues revealed reduced viral replication and altered IFN- and tumor necrosis factor (TNF)-response of these FV3 mutants compared to FV3-WT that were distinct between infected tadpole and adult frogs [22, 23]. Based on the function their host homologs, both viral CARD-like and βHSD mimics are preumably interfering with regulation of host immune responses, particularly inflammatory response [31, 32]. Thus, these viral genes may indirectly intersect to antiviral IFN response through a newly elucidated non-canonical epigenetic regulation [30, 33]. Similarly, the FV3 ICP18 gene, which contains a DUF2888 domain shared within the ranavirus clade, appears to also affect IFN signaling by unknown mechanisms. Therefore, we sought to identify potential ranaviral factors that may directly affect virus-host interaction based on transcriptomic analysis. Through integrative uses of protein domain search, amino acid sequence similarity and structural analyses of hypothetical proteins encoded in the FV3 genome, we identified eight FV3 hypothetical proteins containing regions analogous to interferon regulatory factor (IRF) domain or Fibronectin type 3 (FN3) domain (Table 2). As FN3 is a functional domain in cellular receptors conferring recognition of IFNs and other cytokines, IRF domains of IRF transcription factors are characterized by DNA-binding capacity in the promoters of various IFN-stimulated genes (ISGs) including IFN themselves [34, 35]. Because of the critical role of IFN receptors and IRFs in IFN-mediated antiviral immunity, various antagonisms have been identified [36–38]. Particularly in the NCLDV group that have a large DNA genome, viral mimics counteracting IFN receptors and IRFs have been studied (e.g., human herpesviruses [36–38]). However, no similar ranaviral mimics have been elucidated so far, even though various antagonistic effects on IFN response by FV3 infection have been observed [22,23,28].

As shown in Table 2, most of these viral mimics contain regions resembling one or two IRF domains (vIRFs) of various vertebrate IRF proteins, and two share similarity with FN3 domains (vFN3) of IFN receptor subunits forming the type III IFNs, i.e., IFN-λ receptor 1 (ifnlr1) and interleukin-10 receptor beta (il10rb). In Figure 10 and Figure 11, we performed amino acid similarity alignments of the identified vIRFs with respective IRF domains conserved among vertebrate IRF protein targets. The general IRF consensus comprises a N-terminal DNA-binding domain (DBD) with five typical tryptophan repeats (5W) and a C-terminal activation region (AD) containing an IRF-associated domain (IAD). As DBD is essential for the recognition of DNA motifs within conserved *cis*-regulatory elements (CRE) in the promoter region of IFN or ISG genes, more variable IAD mediates protein-protein interactions with other transcription factors and hence defines functional diversity of different members in the IRF family [34, 35]. Reminiscent of vIRFs identified in human herpesviruses, putative FV3’s vIRFs share a high sequence similarity of positive charged residues and an average ∼30% amino acid sequence identity with DBD and/or AD domains in corresponding amphibian IRFs, but exhibit less similarity with 5W-pattern associated to the DBD domains [34–38]. The identified putative vIRFs exhibit a broad target potential on *Xenopus* IRF1/IRF2 (Norf76R), IRF3 (Norf13L and Orf19R), IRF4 (Orf41R), IRF6 (Norf42L), and IRF8 (Orf27R and Orf82R). In addition to the coding genes of the 98 FV3gorfs (Orf) annotated along the FV3 reference genome, some vIRFs were found encoded by alternative coding frames (Norf), and thus are newly predicted by this study with supportive transcriptomic data. This indicates an extended coding capacity of the FV3 genome beyond our previous understanding (Supplement Excel Sheet). Notably, the temporal class of these viral mimics (except Norfs) has been reported as unknown (UNK) or L in previous studies [17–19], except Orf82R (encoding ICP18) that is an IE gene [19, 23]. This substantiates our previous observation about the increased stimulation of type I and III IFNs in FV3-Δ18K infected tadpole and frog tissues [22, 23]. Studies of human and murine IRFs have shown that IRF4 and IRF8 are highly expressed in lymphoid and myeloid immune cells, and critical for B lymphocyte development and Th cell differentiation. Notably, IRF8 is required for IFN production by dendritic cells (DCs), particularly plasmacytoid DCs (pDCs) that are important IFN producers for early antiviral regulation [25,39,40]. Therefore, ICP18 in ranaviruses is postulated to affect early antiviral IFN responses by targeting amphibian IRF8-mediated IFN responses in lymphocytes including DCs and macrophages [23, 40].

**Figure 10.**
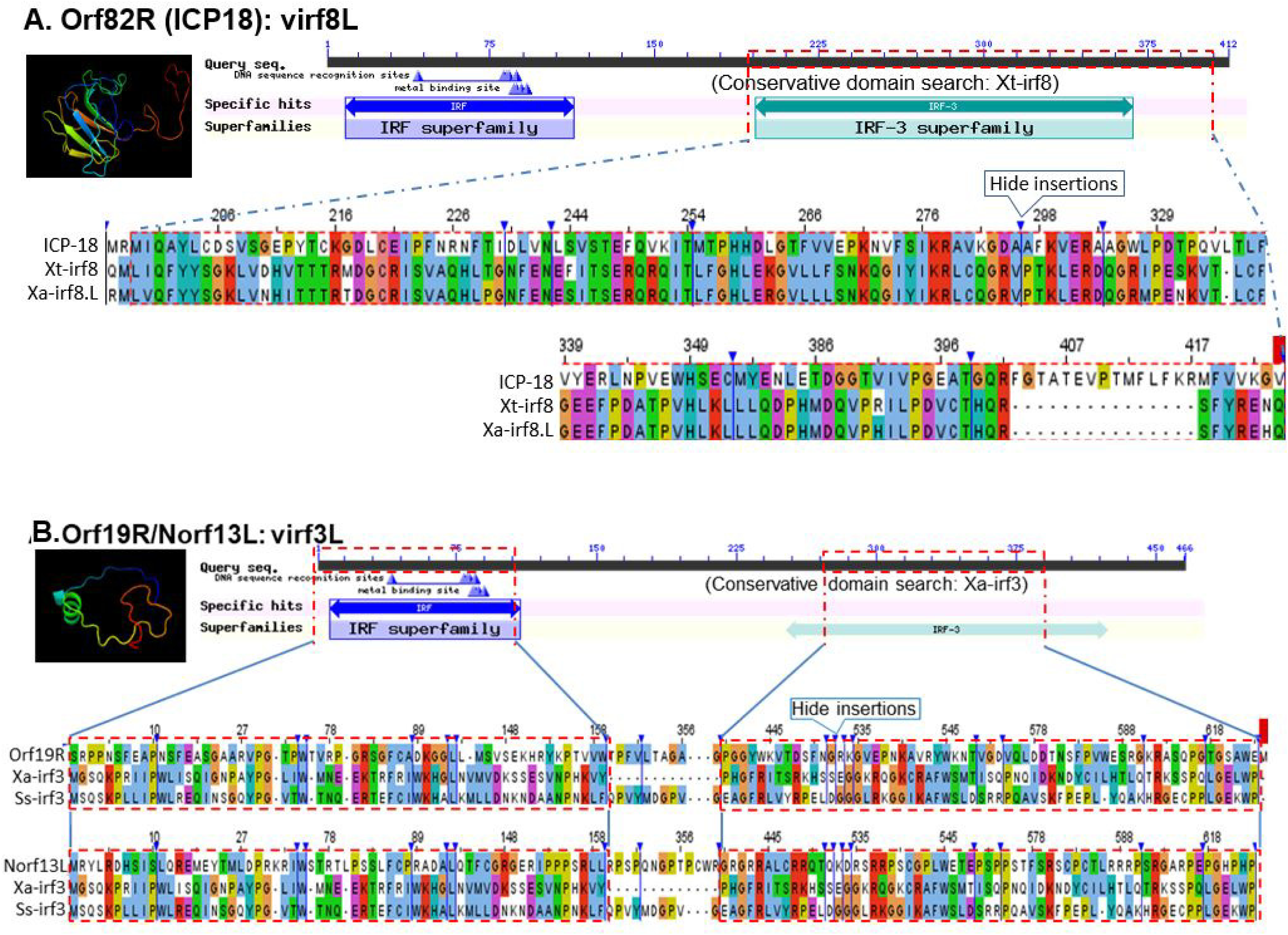
Identification of putative interferon regulatory factor domain (IRF) in FV3 hypothetical proteins, which are potentially virus-coding molecular mimics able to interfere with the host antiviral interferon signaling. (A) Orf82R refers to FV3orf82R spanning 89450-89923 nt region on the positive strand of the FV3 genome (NC_005946.1) and encodes a hypothetical protein ICP-18 of 157 AA. It contains the second IRF-like domains at ∼30% AA sequence similarity to the second IRF domain detected at *Xenopus* IRF8 proteins as illustrated. (B) Orf19R refers to FV3orf19R spanning 21916-24471 nt region on the positive strand of the FV3 genome and encodes a hypothetical protein of 851 AA. It contains two IRF domains (partially second one) at its C-terminal ∼500 AA. Norf13L ascribes a novel open reading frame (Norf) predicted using FGEESV0 program, which spans 14685-15092 nt region of the negative strand of FV3 genome and encodes a hypothetical protein of 135 AA. It contains two nearly consecutive IRF domains (partially second one) as aligned to irf3 homologs. The full-length sequences of the predicted hypothetical ORF/proteins are provided in Supplemental Excel Sheets. The collective information of other IRF-like-domain containing proteins and GenBank Accession Numbers of the aligned protein sequences are listed in Table 2 and diagramed in Figure S2 and S3, respectively. The protein domain analyses were queried and extracted using NCBI CDD database, the tertiary structures were simulated using a Phyre2 and PyMol programs, and sequence alignments and view, with a Jalview program.

**Figure 11.**
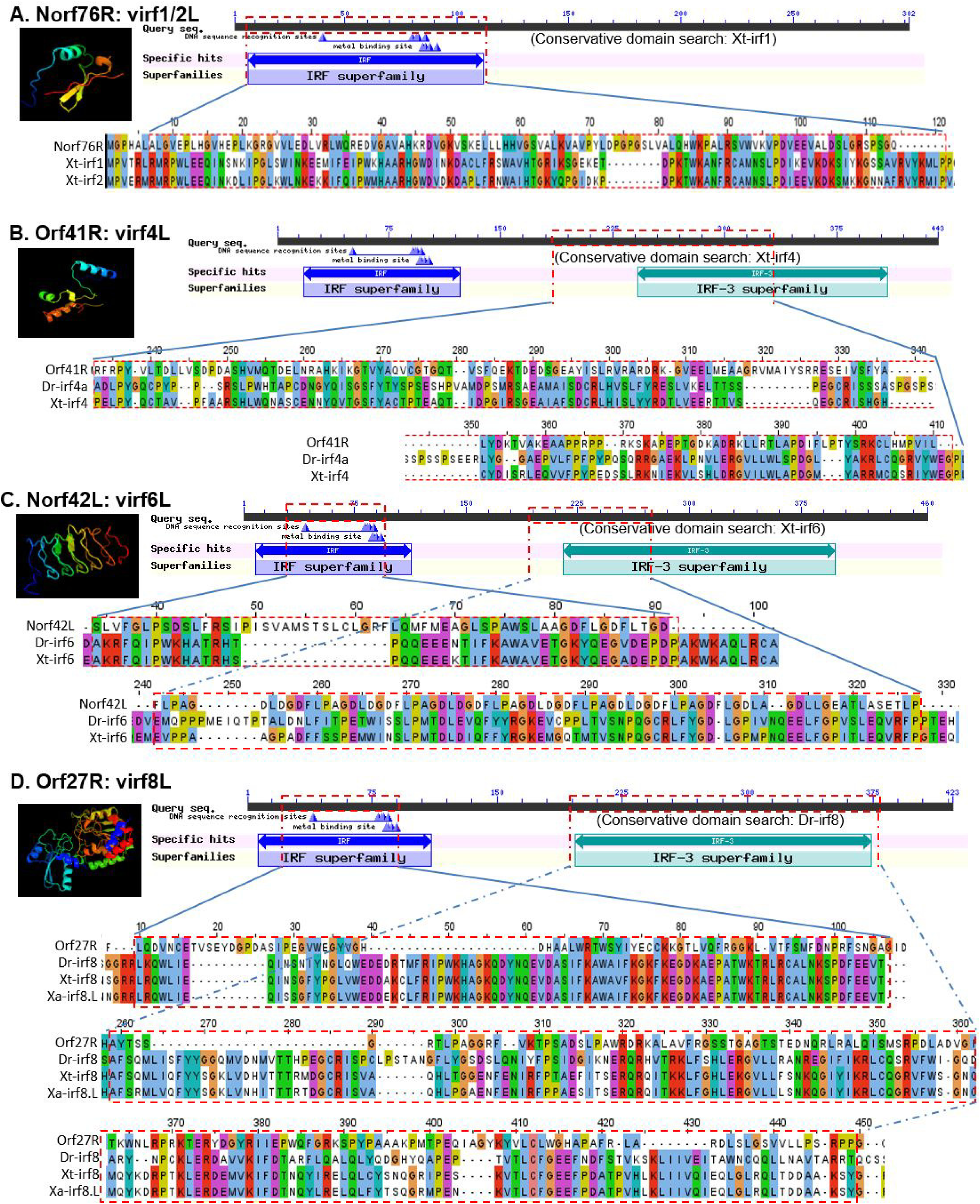
Identification of putative interferon regulatory factor domain (IRF) in several FV3 hypothetical proteins, which are potentially of virus-coding molecular mimics (virfL) to interfere with the host antiviral interferon signaling. (A) Norf76R ascribes a novel open reading frame (ORF) predicted using FGEESV0 program, which spans 66460-67080 nt region of the positive strand of FV3 reference genome (NC_005946.1) and encodes a hypothetical protein at 206 AA. Its N-terminal ∼120 AA contains a domain similar to (∼30% identity) IRF1/2 of *Xenopus tropicalis* (Xt). (B) Orf41R is a consensus to FV3orf41R spanning 46691-50188 nt region on the positive strand of the FV3 genome and encodes a hypothetical protein at 1165 AA. It contains a partial IRF domain similar to Xenopus irf4 at its middle ∼200 AA. (C) Norf42L ascribes a novel predicted ORF, which spans 38635-39102 nt region of the negative strand of FV3 genome and encodes a hypothetical protein at 155 AA. It contains two partial IRF domains in aligned to irf6 homologs. (D) Orf27R is a consensus to FV3orf27R spanning 33728-36640 nt region on the positive strand of the FV3 genome and encodes a hypothetical protein at 970 AA. It contains two IRF domains similar to *Xenopus/Danio* irf8 at its middle ∼400 AA. The full-length sequences of the predicted hypothetical ORF/proteins are provided in Supplemental Excel Sheets. The collection of other IFN-interfering domain containing proteins are listed in Table 2, the programs used for sequence, domain and structure analyses were as in Figure 10.

The main characterized roles of other IRFs include: (1) IRF1 and IRF2 regulate T cell activation and enhance Th1 immune response; (2) IRF3 and IRF7 are engaged in IFN production signaling downstream innate immune recognition of various intracellular pathogens including viruses; (3) IRF5 is involved in the regulation of inflammation and apoptosis, and structurally similar to IRF6 regulates proliferation and differentiation of keratinocytes; and (4) IRF9 is part of the IFN-stimulated gene factor 3 (ISGF3) complex that transmit type I and III IFN signals [35,39,40]. The detection of vIRFs broadly targeting amphibian IRF1/2, IRF3, IRF4, IRF6, and IRF8, is supported by previous and current observations about an IFN-centered and broadly other immune interactions during FV3 infection in amphibians, which also indicates a general cross-species conservation, molecularly and functionally, of these amphibian IRFs in immune regulation (Table 2, Figure 10 and Figure 11) [22,23,28,39].

Figure 12 shows alignments of two vFN3 mimics that contain relevant domain similar to vertebrate interferon lambda receptor 1 (ifnlr1) or IL-10 receptor beta unit (il10rb), which form a functional IFN receptor interacting with type III IFNs in responsive cells [40–43]. As illustrated in this study, Norf66L encodes a novel open reading frame (Norf), which spans a 59162-60037 nt region on the negative strand of the FV3 reference genome and encodes a hypothetical protein at 291 AA. It contains a FN3-like domain (residue 121-230 AA) similar to the ifnlr1 isoforms both in term of sequence similarity and modeled β-sheet containing structure. By contrast, Orf59L refers to FV3orf59L spanning a 65956-67014 nt region on the negative strand of the FV3 genome and encodes a hypothetical protein at 352 AA. It contains a FN3 domain region at 108-203 AA and is molecularly similar to that of *Xenopus* il10rb. The detection of vFN3 mimics that primarily targets type III IFN receptors is consistent with previous observation about FV3’s suppressive effect on IFN-λ expression in *Xenopus* [23, 28], which generally concurs with our host-specific transcriptome analysis (data not shown). It is, therefore, likely that FV3 has strengthened its IFN-antagonism during evolution to overcome the epithelial specific type III IFNs signaling pathway, particularly in tadpoles [23,28,41–43].

**Figure 12.**
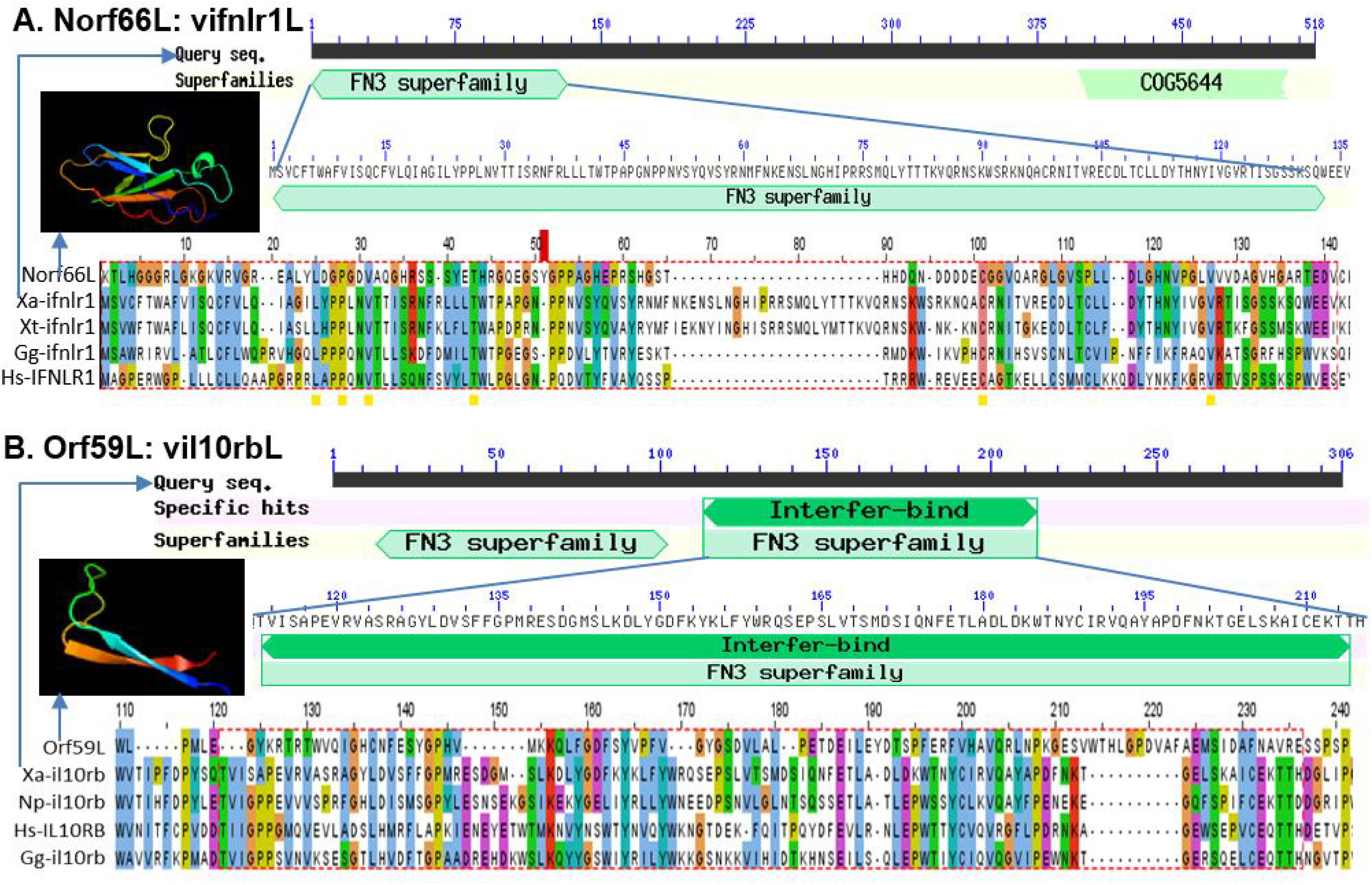
Detection of Fibronectin type 3 domain (FN3) conserved in IFN receptors of several FV3 hypothetical proteins, which are potential virus-coding molecular mimics able to interfere with the host antiviral interferon signaling. (A) Norf66L ascribes a novel open reading frame (Norf), which spans 59162-60037 nt region of the negative strand of FV3 reference genome and encodes a hypothetical protein of 291 AA. It contains a FN3 domain (residue 121-230 AA) similar to vertebrate interferon lambda receptor 1 (ifnlr1) isoforms regarding the sequence similarity and β-sheet containing structure. (B) Orf59L refers to FV3orf59L spanning 65956-67014 nt region on the negative strand of the FV3 genome and encodes a hypothetical protein of 352 AA. It contains a FN3 domain region at 108-203 AA and is molecularly similar to that of *Xenopus* IL-10 receptor beta unit (il10rb). The full-length sequences of the predicted hypothetical ORF/proteins are provided in Supplemental Excel Sheets. The collective information of other IFN-interfering-domain containing proteins and GenBank Accession Numbers of the aligned protein sequences are listed in Table 2 and diagramed in Figure S2 and S3, respectively. The protein domain analyses were queried and extracted using NCBI CDD database, the tertiary structures were simulated using a Phyre2 and PyMol programs, and sequence alignments and view, with a Jalview program.

### Ligand-protein docking analysis for interaction between putative viral mimics and the host factors in IFN signaling

We further performed ligand-protein docking analyses to simulate the DNA-protein interaction between putative vIRFs and respective regulatory *cis*-elements (CRE) extracted from several *Xenopus* IFN gene promoters as determined previously [25]. As shown in Figure 13 and Figure 14, in addition to their sequence similarity to corresponding amphibian IRF domains, these putative vIRFs were docked by respective CREs comparatively to *Xenopus* IRFs despite interfacial difference. Estimated by two interfacial docking parameters, [i.e., the ΔG docking score, which estimates binding affinity energy, and ligand root-mean-square-deviations (Rmsd), which indicates the average distance between the backbone atoms of the ligand to receptors], the Orf60R vs IRF2, Orf41R vs IRF4, and particularly ICP18 vs IRF8, showed very close ΔG and Rmsd values with the other comparative pairs having more difference per either parameters. Notably, the CREs used for IRF docking were extracted from two *X. laevis* IFN genes proximal promoters (*IFN7p* and *IFNLX1p*) [25]. The corresponding CREs from other IFN or ISG gene promoters may conserve these matrix patterns but differ in their exact sequences, which may affect their interaction and docking parameters with vIRFs and amphibian IRFs, thus resulting in gene-specific expressional interference [25].

**Figure 13.**
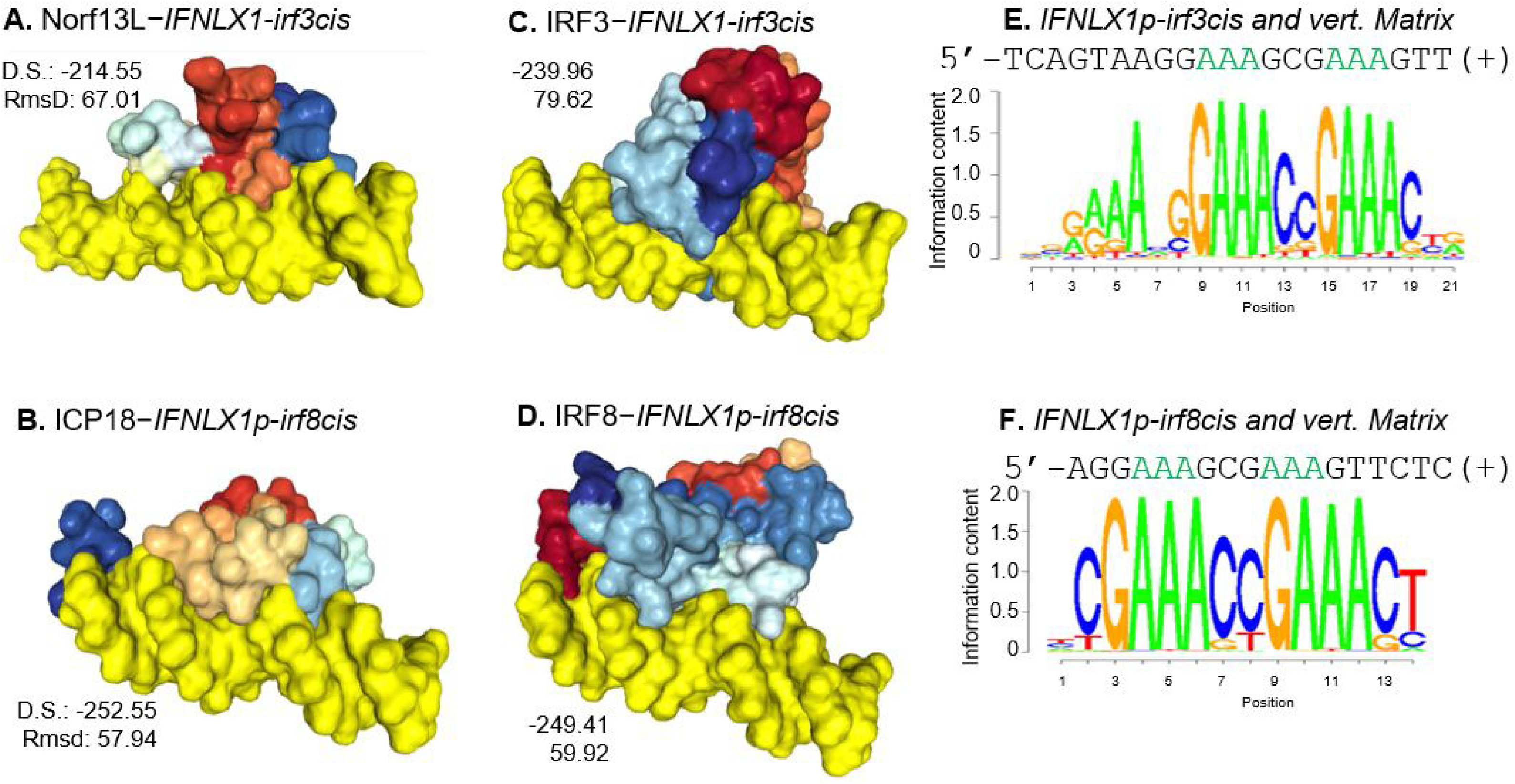
Simulation of protein-DNA docking between putative viral IRF mimics (hypothetical proteins encoded by FV3 ORFs) and respective regulatory *cis*-elements (CRE) extracted from a *Xenopus* IFN gene promoters (IFNLX1p) (Left panel, A and B). The docking structures of respective CREs with corresponding similar *Xenopus* IRFs were performed with same algorithms and defaults for comparison (Mid panel, C and D). Two *X. laevis* IFN genes proximal promoters (*IFN7p* and *IFNLX1p*) were determined as described, and the IRF-binding CREs were determined and extracted using mPWM tools at https://ccg.epfl.ch/pwmtools/pwmscore.php with CRE Matrices are from JASPAR CORE 2020 Vertebrates (vert.) motif library affiliated with the PWM tools (Right panel, K-O). PWM, position weight matrix. Protein-DNA docking and structure analysis was performed similarly as described for Figure 11. Showing also the Docking scores (D.S.), which estimate binding affinity energy (ΔG), and ligand root-mean-square-deviations (Rmsd), which indicate the average distance between the backbone atoms of the ligand to receptors. The full-length sequences of the predicted hypothetical ORF/proteins are provided in Supplemental Excel Sheets. The GenBank Accession Numbers of the aligned protein sequences are in Table 2 and diagramed in Figure S2 and S3, respectively.

**Figure 14.**
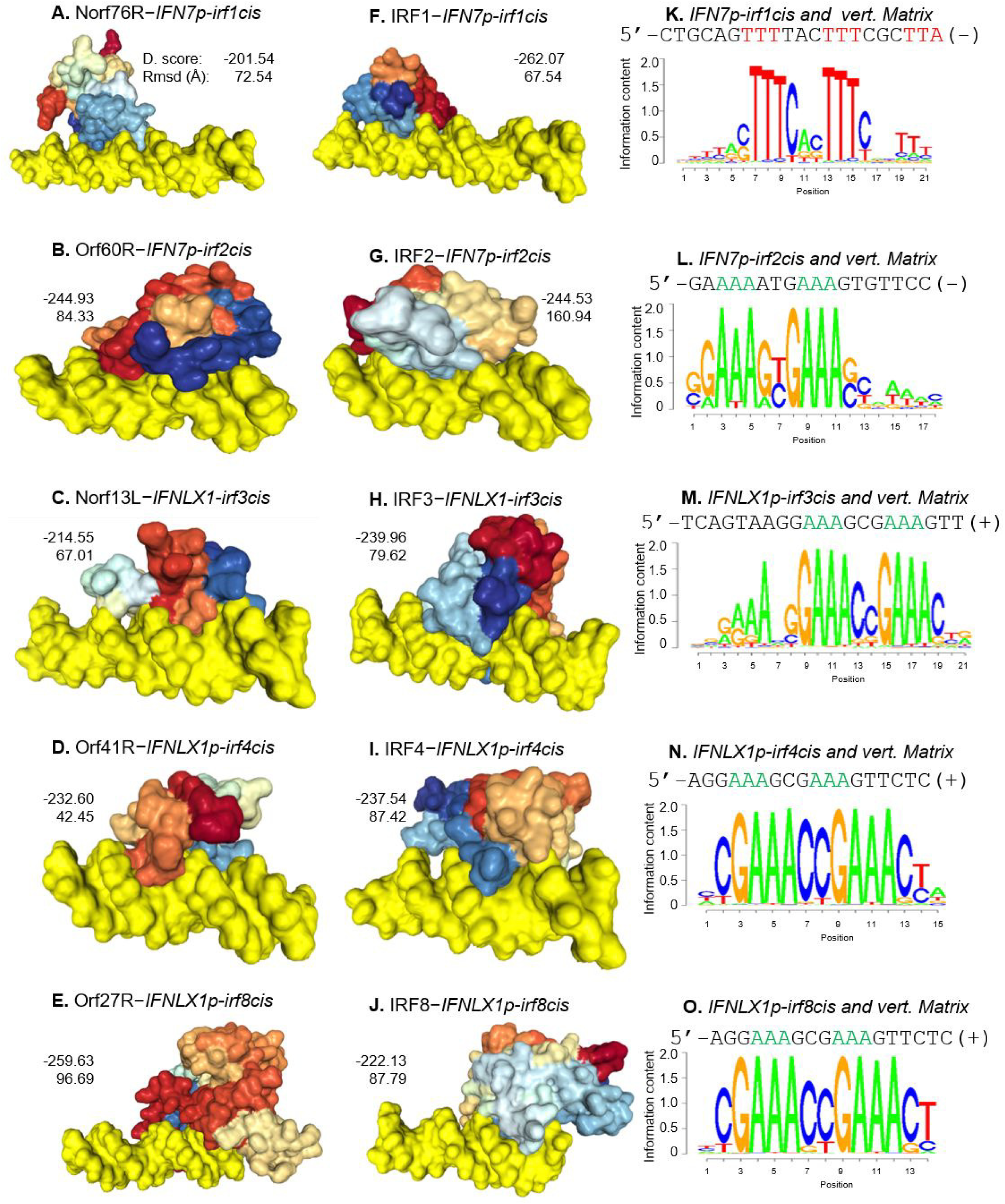
Simulation of protein-DNA docking between putative viral IRF mimics (hypothetical proteins encoded by FV3 ORFs) and respective regulatory *cis*-elements (CRE) extracted from some *Xenopus* IFN gene promoters *(IFN7p* and *IFNLX1p*) (Left panel, A-E). The docking structures of respective CREs with corresponding similar *Xenopus* IRFs were performed with same algorithms and defaults for comparison (Mid panel, G-J). Two *X. laevis* IFN genes proximal promoters (IFN7p and IFNLX1p) were determined as described, and the IRF-binding CREs were determined and extracted using mPWM tools at https://ccg.epfl.ch/pwmtools/pwmscore.php with CRE Matrices are from JASPAR CORE 2020 Vertebrates (vert.) motif library affiliated with the PWM tools (Right panel, K-O). PWM, position weight matrix. Protein-DNA docking and structure analysis was performed similarly as described for Figure 11. Showing also the Docking (D.) scores, which estimate binding affinity energy (ΔG), and ligand root-mean-square-deviations (Rmsd), which indicate the average distance between the backbone atoms of the ligand to receptors. The full-length sequences of the predicted hypothetical ORF/proteins are provided in Supplemental Excel Sheets. The GenBank Accession Numbers of the aligned protein sequences are listed in Table 2.

Similar docking simulation was performed using a *X. laevis* type III IFN ligand (IFN-λ3) to compare between vFN3 mimics and corresponding IFN receptor subunits, IFNLR1 and IL10rb. The two vFN3 mimics, encoded by FV3 Norf66L and Orf59L respectively, were docked by the IFN-λ3 ligand as well as to each of the IFN receptor subunits. However, both vFN3 mimics were docked by IFN-λ3 ligand using different interfacial networks with a higher ΔG (i.e., lower binding affinity) and quite different Rmsd (Figure 15). This indicates that these vFN3 mimics may indirectly act on IFN-ligand competition or induce conformation hindrance rather than directly blocking the IFN-interacting sites on targeted IFN receptor subunits. Thus, these vFN3 likely interfere with the receptor to efficiently detect type III IFN signaling and alter downstream IFN signaling that induces antiviral response [34].

**Figure 15.**
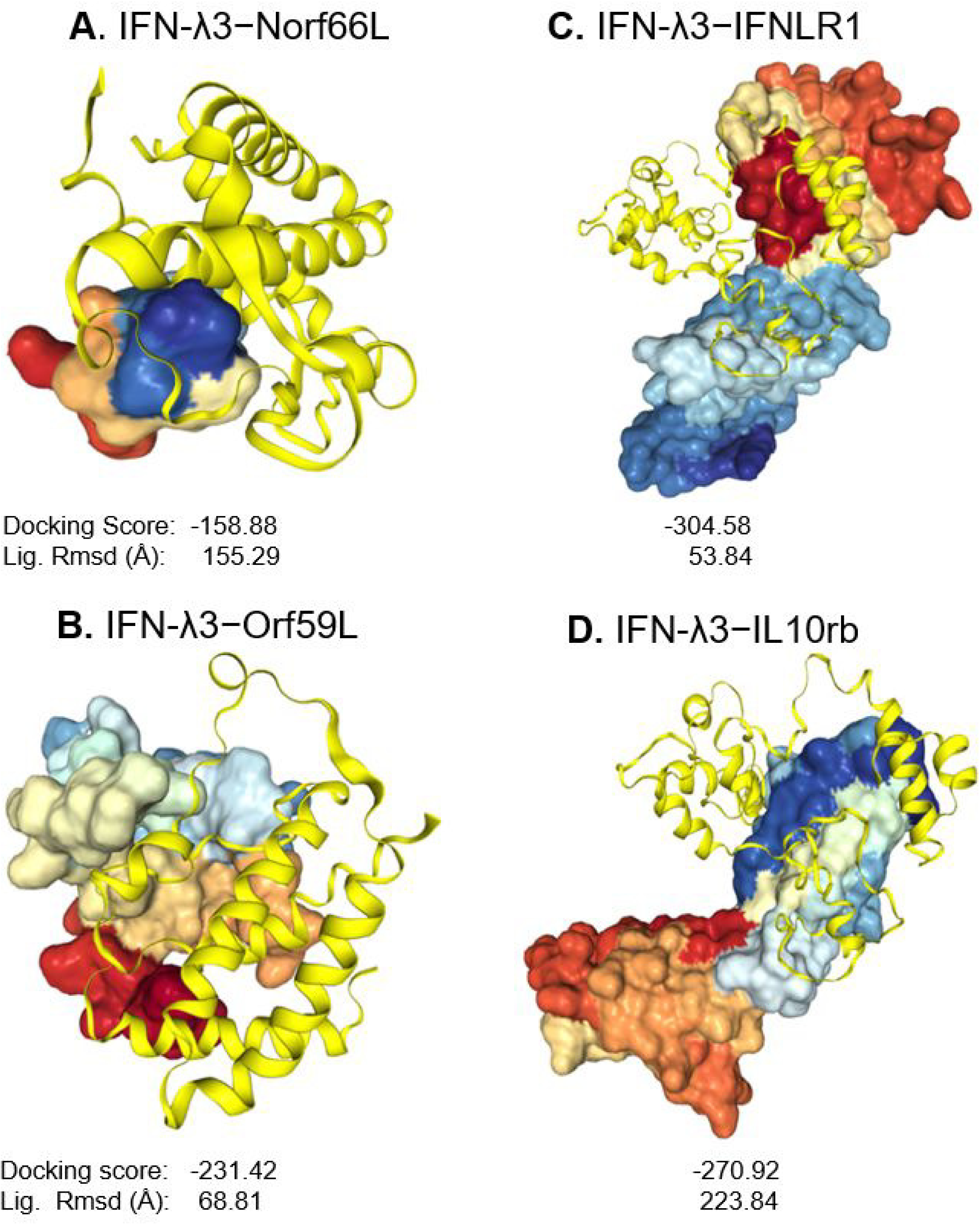
Simulation of protein-protein docking between a X. laevis type III interferon ligand (IFN-λ3, Cartoon Yellow) and two putative viral receptor mimics (Surface energy Rainbow) encoded by two FV3 ORFs Norf66L and Orf59L (Left two, A and B). The docking structures of IFN-λ3 (Cartoon Yellow) with its natural receptor units (IFNLR1 and IL10rb, Surface energy Rainbow) were performed with same algorithms and defaults for comparison (Right two, C and D). Protein structures were simulated and visualized through combinative uses of the programs of PyMol, Chimera, and Phyre2 as described, and primarily a HDOCK server (http://hdock.phys.hust.edu.cn/) for protein-protein docking based on a hybrid algorithm of *ab initio* free docking. Showing also the Docking scores, which estimate binding affinity energy (ΔG), and ligand root-mean-square-deviations (Lig. Rmsd), which indicates the average distance between the backbone atoms of the ligand to receptors. The full-length sequences of the predicted hypothetical ORF/proteins are provided in Supplemental Excel Sheets. The GenBank Accession Number for Xenopus IFN-λ3 is XP_018089340.1, and others of the aligned protein sequences are listed in Table 2.

In summary, based on transcriptome analysis using FV3 infection model in *X. laevis* frogs, we have identified putative ranaviral mimics, vIRFs- and vFN3s-like. The vIRF elements have the potential to interfere with amphibian antiviral immune response, especially by targeting host response mediated by diverse IRF transcription factors ultimately impairing IFN production and action. In addition, vFN3s-like mimics may impair the interaction between type III IFNs with their cellular receptor and attenuate tissue-specific IFN-λ response on epithelial surface, that are critical for FV3 to initiate infection [36–43].

## 4. CONCLUSIVE HIGHLIGHTS

Frog Virus 3 (FV3) represents a well-characterized model to study ranavirus pathogens that are prevalent in worldwide habitats of amphibians, fish and reptiles, and significantly contribute to the catastrophic amphibian decline [5–10]. Based on conventional and novel assignation of FV3 coding genes according to their temporal expression during infection [17–19], the current study used an unbiased transcriptomic RNA-Seq analysis to profile and compare viral transcripts in various tissues of frogs infected with either FV3-WT and a FV3-Δ64R strain defective for a gene encoding a CARD motif. Results revealed a full-genome coverage transcriptome annotated to almost all coding genes at ∼10× depth on both positive and negative strands in RNA samples from infected intestine, liver, spleen, lung and especially kidney. In contrast, only a partial transcript coverage was detected in infected thymus, skin and muscle suggesting inefficient viral replication in these tissues. Extensive analyses indicated a multi-organ infection pattern of FV3 infection in frogs and validated the *in vivo* expression of most annotated 98 ORFs, as well as their differential expression in a tissue-, virus strain-, and temporal class-dependent manners.

About half of FV3’s coding genes have not yet been functionally determined in the scenario of the virus-host interaction. As a first step to fill this gap of knowledge, we focused our transcriptome-initiated functional analyses on putative ORFs that encode hypothetical proteins containing viral mimicking domains such as host interferon (IFN) regulatory factors (IRFs) and IFN receptors. Our findings suggest that ranaviruses like FV3 have acquired during evolution previously unknown molecular mimics interfering with host IFN signaling, which thus provide a mechanistic understanding about ranavirus persistence in the adult frogs. Although this study relying on inferences from transcriptomic and genomic sequence analysis has inherent limitation, it provides an important initial step for more comprehensive functional studies by biochemical characterization and genetic manipulation.

## Author Contributions

Y.T. and F.D.A contributed to conduct experiments, student training, and proof reading; J.R., contributed to conceptualization, funding acquisition, advisory direction, and resource sharing; Y.S. performed overall conceptualization, experimental coordination, data analysis, draft writing, and funding acquisition.

## Acknowledgement

Dr. Yuanying Gong had participated in sampling handling. We appreciate for the input of Dr. Louise A. Rollins-Smith at Vanderbilt University, and Dr. William B. Sutton at Tennessee State University for relevant discussion and funding acquisition.

